# Natural changes in light interact with circadian regulation at promoters to control gene expression in cyanobacteria

**DOI:** 10.1101/193557

**Authors:** Joseph R. Piechura, Kapil Amarnath, Erin K. O’Shea

**Affiliations:** Department of Molecular and Cellular Biology, Harvard University, Cambridge, United States; FAS Center for Systems Biology, Harvard University, Cambridge, United States; Howard Hughes Medical Institute, Harvard University, Cambridge, United States; Department of Chemistry and Chemical Biology, Harvard University, Cambridge, United States

## Abstract

The circadian clock interacts with other regulatory pathways to tune physiology to predictable daily changes and unexpected environmental fluctuations. However, the complexity of circadian clocks in higher organisms has prevented a clear understanding of how natural environmental conditions affect circadian clocks and their physiological outputs. Here, we dissect the interaction between circadian regulation and responses to fluctuating light in the cyanobacterium *Synechococcus elongatus*. We demonstrate that natural changes in light intensity substantially affect the expression of hundreds of circadian-clock-controlled genes, many of which are involved in key steps in metabolism. These changes in expression arise from control of RNA polymerase recruitment to promoters by circadian and light-responsive regulation of a network of transcription factors including RpaA and RpaB. Using phenomenological modeling constrained by our data, we reveal simple principles that underlie the small number of stereotyped responses of dusk circadian genes to changes in light.

---

Circadian clocks allow organisms from almost all branches of life to alter physiology in anticipation of diurnal changes in the environment. Circadian clocks keep time using autonomous core oscillators which control output pathways to direct periodic changes in the mRNA levels (expression) of genes, ultimately leading to oscillations in higher order behaviors [1]. Laboratory studies of the outputs of circadian clocks have been primarily performed under constant conditions to isolate circadian regulation from environmental responses. In nature, however, organisms with circadian clocks must also cope with unexpected fluctuations in the environment. Thus a major challenge in chronobiology is to understand circadian clocks and their outputs in dynamic environments.

Previous studies suggest that circadian clock output pathways interact with environmental responses to tailor physiological outputs to both the time of day and the current state of the environment. For example, sleep/wake cycles in *Drosophila melanogaster* and photosynthesis in *Arabidopsis thaliana* are controlled by both the circadian clock and environmental variables like day length or light [2, 3]. Further, circadian clocks can modulate environmental responses based on the time-of-day in a process called circadian gating [4, 5]. However, the complexity of higher organisms has prevented a detailed understanding of the interaction between circadian timing information and environmental responses. In contrast, the circadian clock in the cyanobacterium *Synechococcus elongatus* PCC7942, an obligate photoautotroph, has a simple architecture which controls gene expression oscillations (Figure 1A) to influence metabolism and growth. *S. elongatus* must carefully monitor its environment, as the sunlight required for photosynthesis fluctuates on the minute, day, and seasonal timescales (Figure 1B, [6]). While it is well understood how the circadian clock in *S. elongatus* behaves under constant conditions, it is unclear how this system changes in natural, fluctuating light.

Under ‘Constant Light’ conditions (Figure 1A, dashed navy blue line), genes which show oscillatory expression (circadian genes) can be divided into two groups, the dawn and the dusk genes, which peak at subjective dawn and subjective dusk [7, 8] (Figure 1A). Subjective dawn and subjective dusk refer to the times at which dark-to-light or light-to-dark transitions would occur in a 12 hr light-12 hr dark environmental cycle. The dawn genes consist of the core metabolic and growth genes for *S. elongatus*, including the photosystems, ATP synthase, carbon fixation/Calvin-Benson-Bassham cycle enzymes and ribosomal proteins [7–9]. In the absence of regulation by the circadian clock under Constant Light, *S. elongatus* constantly expresses dawn genes [10]. The clock primarily regulates the expression of dusk genes [10], which include the genes required to utilize glycogen as an energy source in the absence of sunlight, such as glycogen phosphorylase and cytochrome c oxidase. As such, the circadian clock serves a critical function in switching *S. elongatus* from a daytime state of photosynthesis to a nighttime state of carbon metabolism through glycogen breakdown [9, 11–13]. In Constant Light conditions, the dusk and dawn genes show oscillatory expression with a 24 hr period, resulting in broad peaks of maximal expression (Figure 1A, solid green line and dashed red line) [7, 8]. Recent whole-cell modeling of metabolism, protein levels, and growth predict that this picture of circadian gene expression should change under the dynamic light conditions of a natural, clear day (Figure 1B, navy blue line) [14]. The modeling suggests that making and using glycogen is a major cost to cell growth and thus the expression of genes required to switch metabolism from photosynthesis to glycogen breakdown should be delayed until absolutely necessary [14]. However, gene expression in natural light conditions has not been measured in *S. elongatus*.

Consistent with predictions of light-dependent changes in circadian gene expression, current evidence suggests interaction between circadian and light regulatory pathways. The cyanobacterial clock keeps track of the time of day using a core post-translational oscillator (PTO) that consists of three proteins, KaiA, KaiB, and KaiC, whose enzymatic activities result in 24 hr oscillations in the phosphorylation state of KaiC [15–17]. In vivo under Constant Light conditions the Kai PTO modulates circadian gene expression by controlling oscillations in phosphorylation state of the master OmpR-type transcription factor RpaA [10, 18] to peak at subjective dusk (Figure 1A, black dotted line; Figure 1C) [18, 19]. Phosphorylated RpaA (RpaA∼P) binds to the promoters of some dusk genes to activate their expression, leading to indirect activation of other dusk genes and repression of dawn genes (Figure 1C) [10]. As *kaiBC* is a dusk gene target of RpaA, the Kai PTO directs its own expression, resulting in a transcription-translation feedback loop (TTL) that stabilizes the phase of the clock [20–22]. Exposure to complete darkness at specific times of day causes phase shifts in the PTO to align clock output with the external day/night cycle [23], in a process called entrainment. However, it is not understood whether RpaA activity and the TTL loop change in the presence of more subtle natural light changes during the day (Figure 1C). Meanwhile the OmpR-type transcription factor RpaB binds to some circadian gene promoters [24], and the phosphorylation state and DNA binding activity of this protein decreases in response to high light exposure (Figure 1C) [25, 26]. However, it is not clear how natural light changes like sunset or shade pulses affect RpaB activity (Figure 1C). Moreover, RpaB clearly plays some role in altering circadian gene expression in response to light[27], but it is unclear how (Figure 1C). While light likely exerts global, growth-rate-dependent regulation [28–30], the interaction between circadian and light regulation to control the activities of RpaA and RpaB represents a particularly tractable scenario for dissecting the mechanisms underlying interaction between clock and environment to control physiology.

Here we measure and model circadian gene expression and several layers of regulation in cyanobacteria grown under the fluctuating light intensities typically experienced in nature. We find that fluctuations in light alter the expression patterns of almost all circadian genes. We identify key regulatory steps at which information about changes in light interact with clock output pathways to control gene expression, and reveal a complex regulatory network underlying circadian gene expression in natural conditions. Finally, we show that phenomenological models effectively describe the integration of the circadian clock with responses to environmental fluctuations.

## Results

### A. Sunlight on a clear day delays the timing of circadian gene expression relative to constant light conditions

To grow and assay cyanobacteria in natural light conditions, we custom-built a culturing setup with a light source that can be programmed to mimic natural fluctuations in sunlight. On a cloudless ‘Clear Day,’ light intensity varies in a parabolic manner due to the rotation of the Earth, ending with a gradual ramp down of light intensity prior to dusk (‘Sunset’, Figure 1B). Rapid changes in cloud cover cause abrupt increases (‘High Light pulse’) and decreases (‘Shade pulse’) in sunlight [6] (Figure 1B). Using a set of programmable warm white LED arrays (Materials and Methods, Construction of light apparatus and Calibrating light conditions) for illumination, in all experiments we grew cells in either a Clear Day that peaked at 600 *µ*mol photons m^−2^ s^−1^ or a constant Low Light of 50 *µ*mol photons m^−2^ s^−1^ (Figure 2A, top panel) for at least two days to acclimate and synchronize the cells before measurement.

To determine whether a natural light profile affects circadian output, we compared genome-wide gene expression in Clear Day conditions versus Low Light conditions using RNA sequencing (Figure 2A, Setup, arrows indicate sampling). We maintained cultures in their respective condition followed by 12 hours of darkness for 3 light/dark cycles, and sampled them (arrows) over the third light period (Figure 2A, Setup). Growth in Clear Day conditions affected pigment levels and growth rate compared to Low light conditions (Figure 2-figure supplement 1). We focused our analysis on a set of high amplitude circadian genes that show oscillatory expression under Constant Light conditions (Figure 2-figure supplement 2; see Materials and Methods, Definition of circadian genes). The Low Light condition (Figure 2B, upper panel) reproduces the expression profile previously observed under Constant Light conditions (Figure 2-figure supplement 2). However, in the Clear Day condition 148 of the 281 dusk genes were expressed two fold or higher at dusk compared to Low Light, demonstrating light-dependent expression. Dawn genes show the opposite behavior — they have higher expression at midday under Clear Day conditions, although this trend is less pronounced (Figure 2-figure supplement 3). Taken together, Clear Day conditions significantly influence the expression dynamics of almost all circadian genes, with the strongest effects on dusk genes.

To look more closely at how the Clear Day condition affects the dusk genes, which are the primary regulatory targets of the clock, we analyze the gene expression dynamics of the representative dusk gene *Synpcc7942 1567*. Under Low Light conditions, *Synpcc7942 1567* exhibits an increase in expression from dawn to dusk, reaching a plateau by 8 hours after dawn (Figure 2C, black solid line). Under Clear Day conditions, however, the expression of this gene remains low through the midday peak of light intensity (Figure 2C, magenta solid line; 4-8 hours after dawn), and its expression sharply increases just prior to dusk as light intensity decreases, reaching maximal expression just as the 12 hour dark period begins. This delayed pattern of gene expression can be seen in almost all dusk genes (Figure 2B; *Synpcc7942 1567* indicated with arrows). Thus Clear Day conditions significantly alter the dynamics and amplitude of dusk gene expression to peak just before dusk.

**Figure 1:**
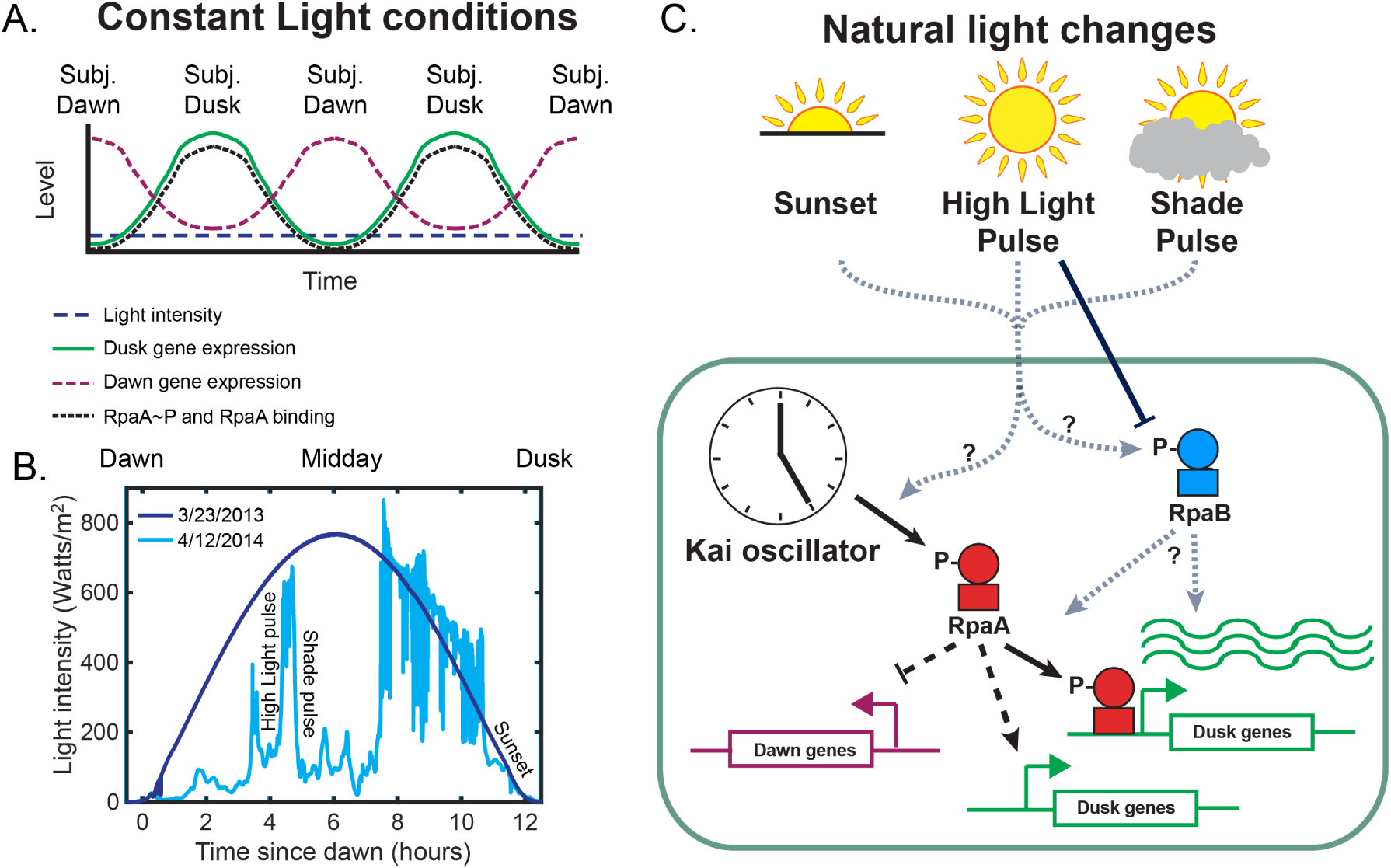
The circadian and light response pathways in cyanobacteria. **(A)** Schematic of gene expression output of the circadian clock under Constant Light conditions. Under Constant Light intensity (blue dashed line), dawn gene expression (magenta dashed line) and dusk gene expression (solid green line) display oscillatory patterns, peaking at subjective dawn and subjective dusk, respectively. The Kai PTO generates oscillations in the levels of phosphorylated RpaA (RpaA∼P) and the binding of RpaA to DNA (black dotted line), with the peak amplitude at subjective dusk. **(B)** Solar irradiance measurements in units of watts m^−^2 at 342.5 meters above sea level in Madison, WI on a clear day (3/23/13, dark blue), and a day on which fluctuations in cloud cover generated rapid changes in light intensity (4/12/14, light blue) [6]. Examples of a ‘High Light pulse,’ ‘Shade pulse,’ and ‘Sunset’ are indicated. **(C)** Schematic of the regulation of circadian gene expression. RpaA phosphorylation state converts timing information from the Kai oscillator to changes in gene expression by directly binding and activating a subset of dusk genes, indirectly activating the remainder of the dusk genes, and indirectly repressing the dawn genes. It is unclear whether natural fluctuations in light directly affect the clock and its output pathways and how light-induced changes in RpaB activity might be involved.

The delay of dusk gene expression likely enables cyanobacteria to switch to inefficient energy production by glycogen breakdown only when absolutely necessary so that they can survive the extended darkness of night. The two glycogen breakdown genes, *glgP* and *glgX*, are both light-dependent dusk genes that strongly peak in Clear Day at dusk, while *glgC*, which codes for the rate limiting enzyme of glycogen synthesis, is a dawn gene whose expression is higher in Clear Day conditions compared to Low Light (Figure 2-figure supplement 4).

These gene expression dynamics would favor both the maintenance of glycogen synthesis until the end of the day and a delay in the activation of glycogen breakdown until just before it is required at nighttime, in agreement with predictions from metabolic modeling during the same Clear Day conditions used here [14]. Thus, environmental conditions are integrated into the output of the circadian clock to potentially optimize resource allocation in naturally-relevant diurnal cycles, as recently suggested [14].

Remarkably, though in both light conditions the cells experience 50 *µ*mol photons m^−2^ s^−1^ at the end of the day just before night, light-dependent dusk genes have substantially higher expression in the Clear Day conditions relative to the Low Light conditions (Figure 2B-C). This strong activation of dusk genes occurs concomitant with the decrease in light intensity during Clear Day that mimics Sunset, which suggests that *changes* in light intensity affect activation of dusk genes, as opposed to absolute light intensity levels. Dusk gene expression could thus happen ‘just-in-time’ before night regardless of when Sunset occurs.

### B. Changes in light intensity control the transcription of circadian genes

We next asked whether changes in light intensity are a factor controlling the expression of circadian genes. To directly probe how changes in light affect dusk gene expression, we exposed cells to a High Light pulse or a Shade pulse and measured genome-wide gene expression using RNA sequencing. We grew cultures in either Low Light or Clear Day conditions for three days (Figure 3A-B, Setup). On the fourth day, 8 hours after dawn, when RpaA is most active, we exposed the cells to a High Light pulse (Figure 3A) or a Shade pulse (Figure 3B) for 1 hour before returning to the original condition, and sampled them before, during, and after the perturbation. The expression of dusk genes rapidly changed in a direction opposite to the change in light intensity (Figure 3C, all dusk genes; Figure 3E, example dusk gene; Figure 3D, all dusk genes; Figure 3F, example dusk gene), as expected from the effects of the decrease in light intensity at Sunset of the Clear Day condition on circadian gene expression (Figure 2D-E). When cultures were restored to their original condition (High Light to Low Light, Figure 3C,E; Shade to Clear Day, Figure 3D,F), dusk gene expression quickly reverted to a level comparable to that before the pulse. Thus, light-induced changes in dusk gene expression are reversible and responsive to successive shifts in light availability. Dawn gene expression showed the opposite behavior of dusk genes, albeit with less dramatic changes (Figure 3-figure supplement 1). Hence, decreases in light intensity favor the expression of dusk genes (Sunset in Clear Day, Figure 2; Clear Day to Shade and High Light to Low Light, Figure 3), while increases in light favor the expression of dawn genes (midday peak in Clear Day, Figure 2-figure supplement 2; Shade to Clear Day and Low Light to High Light, Figure 3-figure supplement1). Given the more substantial effects of light on dusk gene expression, we focus our further analysis on these genes. To cause these reversible changes in the mRNA levels of dusk genes, changes in light intensity must affect the either the transcription or degradation of dusk gene mRNAs.

To determine whether changes in light intensity regulate the recruitment of RNA polymerase (RNAP) to dusk gene promoters, we performed chromatin immunoprecipitation followed by high-throughput sequencing (ChIPseq) of RNAP in cells immediately before the High Light or Shade pulse (8 hours after dawn in Low Light or Clear Day), and then again after 15 or 60 minutes during the pulse. RNAP enrichment decreased upstream of dusk genes after exposure to High Light, while RNAP enrichment increased after exposure to Shade, correlating with changes in downstream dusk gene expression (Figure 2G,H; Figure 2-figure supplement 2). Thus, changes in light affect RNAP recruitment to dusk gene promoters, suggesting that light conditions affect dusk gene mRNA levels primarily by regulating rates of transcription. These results point to a potential interaction between sunlight and signaling pathways upstream of RNAP. We next explored how the observed changes in dusk gene expression in the presence of natural light fluctuations (Figures 2,3) could be achieved via gene regulatory mechanisms.

### C. Regulation of dusk gene expression by RpaA and RpaB under dynamic light regimes

We first assessed whether changes in light intensity affect RpaA activity to regulate the recruitment of RNAP to dusk genes. Levels of RpaA∼P increased from dawn to dusk similarly in cells grown in either Low Light or Clear Day conditions, and abrupt changes in light intensity did not affect these dynamics. (Figure 4A,B; Figure 4-figure supplement 1). Thus, these natural light fluctuations do not affect the phase of the Kai PTO nor the control of RpaA∼P levels by the Kai PTO (Figure 4E), and thus do not serve as entrainment signals to the clock. However, ChIP-seq showed that light intensity fluctuations alter RpaA∼P binding upstream of dusk genes (Figure 4C; Figure 4-figure supplement 2) in conjunction with RNAP binding upstream of the same gene (Figure 4D; Figure 4-figure supplement 3). The binding of RpaA∼P and RNAP correlated with changes in downstream dusk gene expression (Figure 4C; Figure 4-figure supplement 2). Thus, light fluctuations control the binding of RpaA∼P and RNAP to promoters, suggesting that light-induced changes in the binding of these factors may modulate the activation of dusk gene expression (Figure 4E).

Interestingly, RpaA regulation at a small number (∼10) of promoters including that of *kaiBC* is not substantially affected by light intensity (Figure 4C,D - points around origin; Figure 4-figure supplement 4), demonstrating that the light-dependent regulation of RpaA binding is locus-specific. The KaiABC clock regulates RpaA∼P levels independent of changes in light intensity (Figure 4A,B), and *kaiBC* gene expression dynamics do not substantially change in the Clear Day conditions compared to the Low Light condition (Figure 4-figure supplement 4H-K). Hence, the stabilizing PTO/TTL circadian circuit is robust to natural fluctuations in sunlight. The circadian clock can thus control gene expression independent of environmental changes at select promoters. It is possible that RpaA∼P binding to some promoters is dependent on the association of RNAP with that promoter. As such, regulation that affects RNAP binding to a specific promoter, such as that by sigma factor activity [31], could affect RpaA∼P binding to select promoters.

We next asked whether RpaB plays a role in controlling light-dependent expression of circadian genes. We observed that levels of RpaB∼P changed rapidly in a direction opposite to the change in light (Figure 5A,B; Figure 5-figure supplement 1), suggesting that light affects RpaB activity through its phosphorylation state (Figure 5E). Using ChIP-seq, we found that RpaB binds upstream of a large subset of dusk genes (42/281 dusk genes, Figure 5-source data 2). RpaB binding upstream of these genes shifts after rapid changes in light (Figure 5C; Figure 5-figure supplement 2), correlating with changes in RpaB∼P levels (Figure 5A,B), RNAP binding upstream of the same gene (Figure 5D; Figure 5-figure supplement3), and downstream dusk gene expression (Figure 5C; Figure 5-figure supplement 2). These results suggest that RpaB∼P directly activates the expression of many dusk genes by binding to promoters with RNAP (Figure 5E). Thus, changes in sunlight can regulate dusk genes by adjusting RpaB∼P levels (Figure 5E).

**Figure 2:**
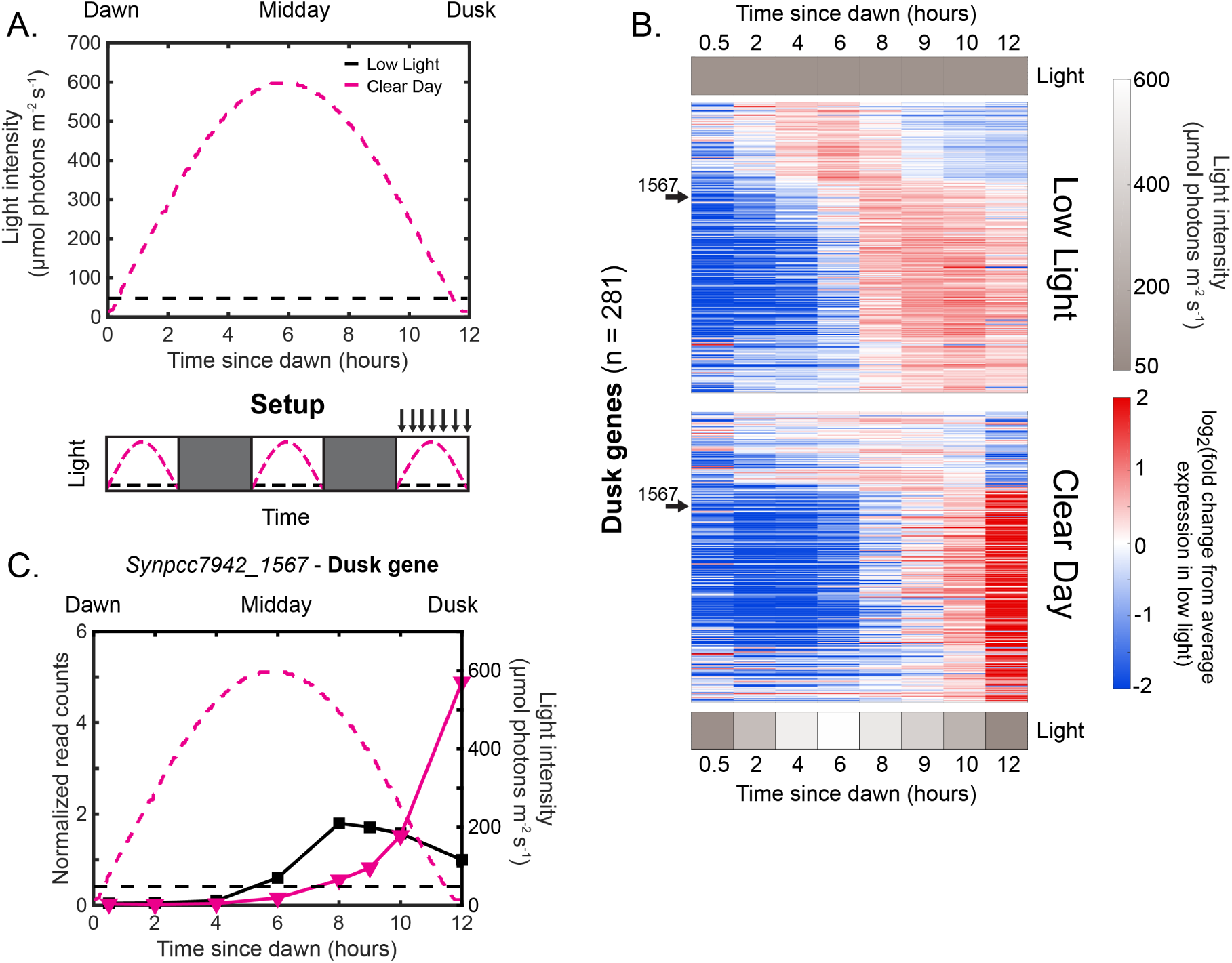
Natural clear day conditions sharpen the expression of dusk genes to peak just before expected darkness. **(A)** Experimental setup for testing the effects of Clear Day conditions on circadian gene expression. The upper plot shows the light intensity profiles of Low Light (black) and Clear Day (magenta) conditions, in units of *µ*mol photons m^−2^s^−1^. The lower plot displays the experimental setup. Cells were grown under Clear Day (magenta dashed lines) or Low Light conditions (black dashed lines), followed by 12 hours of darkness (dark gray boxes) for three days, with period sampling over the third light period (indicated by arrows above plot). **(B)** Gene expression dynamics of all dusk genes (*n* = 281) under Low Light (top) and Clear Day (bottom) conditions. Gene expression is quantified as the log_2_ fold change from the average expression of the gene over all time points in the Low Light condition (see Materials and Methods - RNA sequencing for more details; data available in Figure 2-source data 1). Genes were sorted by phase under Constant Light conditions [8]. Light intensity at each time point is indicated in a grayscale heat map adjacent to the corresponding condition. The data for a representative dusk gene, *Synpcc7942 1567*, is indicated with arrows. **(C)** Gene expression dynamics of the representative dusk gene *Synpcc7942 1567* under Low Light (black) and Clear Day (magenta) conditions (left y-axis). The light profile for each condition is plotted as dashed lines of the same color with values corresponding to the right y-axis. figure supplement 1 (p. 21). Pigment levels of cyanobacteria grown under Low Light or Clear Day conditions indicate adjustments in the photosynthetic apparatus to optimize growth in different light conditions. figure supplement 2 (p. 22). Gene expression dynamics of dusk and dawn circadian genes under Constant Light conditions. figure supplement 3 (p. 23). Dawn gene expression increases during the early part of Clear Day relative to Low Light conditions. figure supplement 4 (p. 24). The gene expression dynamics of glycogen production and breakdown enzymes are changed under Clear Day conditions. source data 1. Normalized gene expression in Low Light and Clear Day conditions.

**Figure 3:**
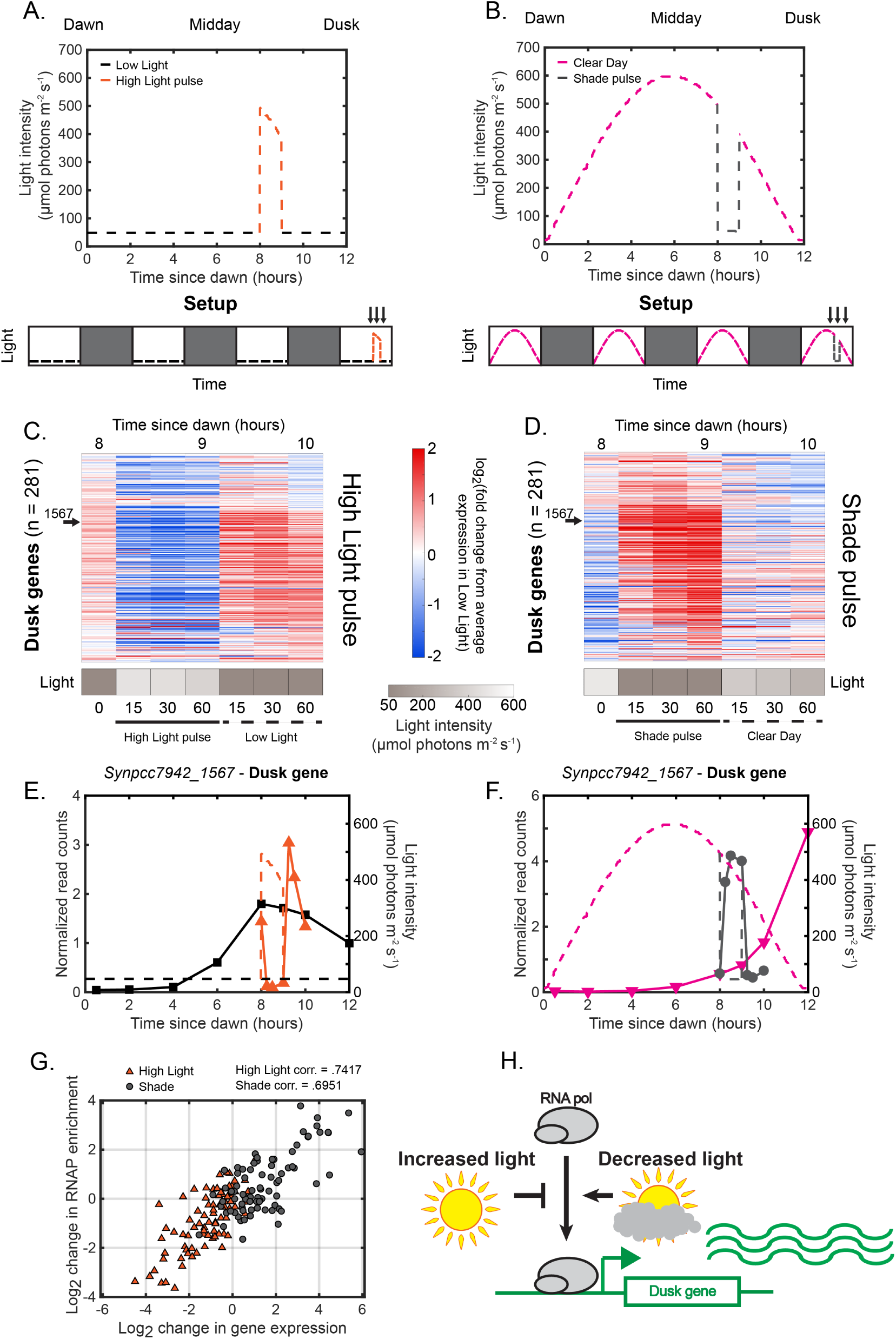
Rapid changes in light intensity modulate the recruitment of RNA polymerase to dusk genes to control dusk gene expression. **(A)** Light intensity profiles of Low Light (black) and High Light pulse (orange) conditions, in units of *µ*mol photons m^−2^ s^−1^. Experimental setup is displayed in the lower plot. Cells were grown for 12 hours under Low Light conditions (black dashed lines), followed by 12 hours of darkness (dark gray boxes) for three days, and then exposed to a High Light pulse (orange dashed lines) at 8 hours after dawn during the fourth light period for one hour before being returned to Low Light conditions. Cells were sampled immediately before, during, and after the High Light pulse (indicated by arrows above plot). (Caption continued on next page.) Figure 3: **(B)** Light intensity profiles of Clear Day (magenta) and Shade pulse (gray) conditions, in units of *µ*mol photons m^−2^ s^−1^. Experimental setup is displayed in the lower plot. Cells were grown for 12 hours under Clear Day conditions (magenta dashed lines), followed by 12 hours of darkness (dark gray boxes) for three days, and then exposed to a Shade pulse (orange dashed lines) at 8 hours after dawn during the fourth light period for one hour before being returned to Low Light conditions. Cells were sampled immediately before, during, and after the High Light pulse (indicated by arrows above plot). **(C)** Gene expression dynamics of dusk genes (*n* = 281) under High Light pulse conditions. Gene expression is quantified as the log_2_ fold change from the average expression of the gene over all time points in the Low Light condition (see Materials and Methods, RNA sequencing for more details; data available in Figure 3-source data 1). Light intensity at each time point in the High Light pulse condition is indicated in a grayscale heat map adjacent to the corresponding time point. **(D)** Gene expression dynamics of dusk genes (*n* = 281) under Shade pulse conditions, plotted as in (C). Genes are ordered the same in **(C)** and(D), sorted by phase under Constant Light conditions [8]. Data for the representative dusk gene *Synpcc7942 1567* is indicated by arrows in **(C)** and (D). **(E)** Gene expression dynamics of the representative dusk gene *Synpcc7942 1567* under Low Light (black) and High Light pulse (orange) conditions (left y-axis). The light profile for each condition is plotted as dashed lines of the same color with values corresponding to the right y-axis. **(F)** Gene expression dynamics of the representative dusk gene *Synpcc7942 1567* under Clear Day (magenta) and Shade pulse (gray) conditions, plotted as in (E). **(G)** Correlation between change in dusk gene expression and the change in enrichment of RNAP upstream of that gene after rapid changes in light intensity. The change in gene expression of a dusk gene (x-axis) and the corresponding change in RNAP enrichment upstream of that gene (y-axis) from the original condition after 60 min in High Light (orange triangles) or Shade (gray circles), plotted for the 82 dusk genes with detectable RNAP peaks in their promoters. See Materials and Methods, ChIP-seq analysis for more details. Data is available in Figure 3-source data 2. The correlation coefficient between change in RNAP enrichment and change in downstream gene expression for the High Light and Shade conditions is indicated above the plot. **(H)** Regulation of RNAP recruitment to dusk genes by changes in light intensity. Increases in light intensity tend to repress the recruitment of RNAP to dusk genes to repress their expression (High Light pulse, Clear Day - midday), while decreases in light intensity (Shade pulse, Sunset of the Clear Day) tend to promote the recruitment of RNAP to dusk genes to activate their expression. figure supplement 1 (p. 25). Rapid changes in light intensity affect dawn gene expression in an opposite direction compared to dusk gene expression. figure supplement 2 (p. 26). Changes in RNAP enrichment and downstream dusk gene expression after rapid changes in light intensity. source data 1. Normalized gene expression in High Light pulse and Shade pulse conditions. source data 2. List of RNAP peaks, gene targets, and quantification of enrichment under High Light pulse and Shade pulse conditions.

Because RpaA and RpaB bind only a subset of light-responsive dusk genes (Figure 6A,B), additional factors must be involved in controlling light-responsive dusk gene expression. Sigma factors are sequence-specific RNAP subunits which regulate gene expression in bacteria [31]. Interestingly, RpaA, RpaB, and RNAP bind to the promoters of three sigma factor genes after abrupt changes in light intensity, correlating with light-responsive changes in expression of these genes (Figure 6C,D; Figure 6-figure supplement 1A-F). These sigma factor genes show light-dependent dusk gene expression patterns (Figure 6-figure supplement 1G-L) that mirror those of the larger group of dusk genes (Figures 2,3), suggesting that these sigma factors could regulate the expression of other dusk genes. Thus, RpaA and RpaB may indirectly regulate the expression of non-target dusk genes by controlling the circadian and light-responsive expression of sigma factor genes [24], similar to how RpaA drives all dusk gene expression in Constant Light conditions by binding to a subset of dusk genes [10]. Given that secondary transcriptional regulators like sigma factors can be regulated post-transcriptionally [31], it is also possible that changes in light intensity affect the activity of dusk transcriptional regulators in a manner independent of RpaA∼P and RpaB∼P regulation. Further, global growth-rate-dependent gene regulatory mechanisms [32, 33] likely cause some of the light-dependent changes in circadian gene expression due to unavoidable differences in the growth rate in different light conditions (Figure 2-figure supplement 1).

We have defined a regulatory picture in which changes in light intensity affect the activity of RpaA and RpaB to control the expression of dusk genes. However, light affects RpaA activity in complex and promoter-specific ways. Additionally, light-dependent regulation in addition to that mediated by RpaA and RpaB may control dusk gene expression in response to environmental perturbations. Still, despite the apparent complexity of regulation of dusk genes in response to light fluctuations, the expression of almost all dusk genes show strikingly regular dynamics (Figure 2,3). Further, the activity of RpaA and RpaB at a subset of promoters (especially those of sigma factor genes) could lead to pervasive and coordinated changes in the expression of other dusk genes. Hence, we reasoned that mathematical models [34] of RpaA and RpaB activity might effectively describe the regulatory circuits underlying the dynamics of large groups of dusk genes. Such an approach would enable an understanding of the basic principles of interaction between circadian gene expression regulation with light-dependent regulation without needing to describe all underlying molecular mechanisms.

**Figure 4:**
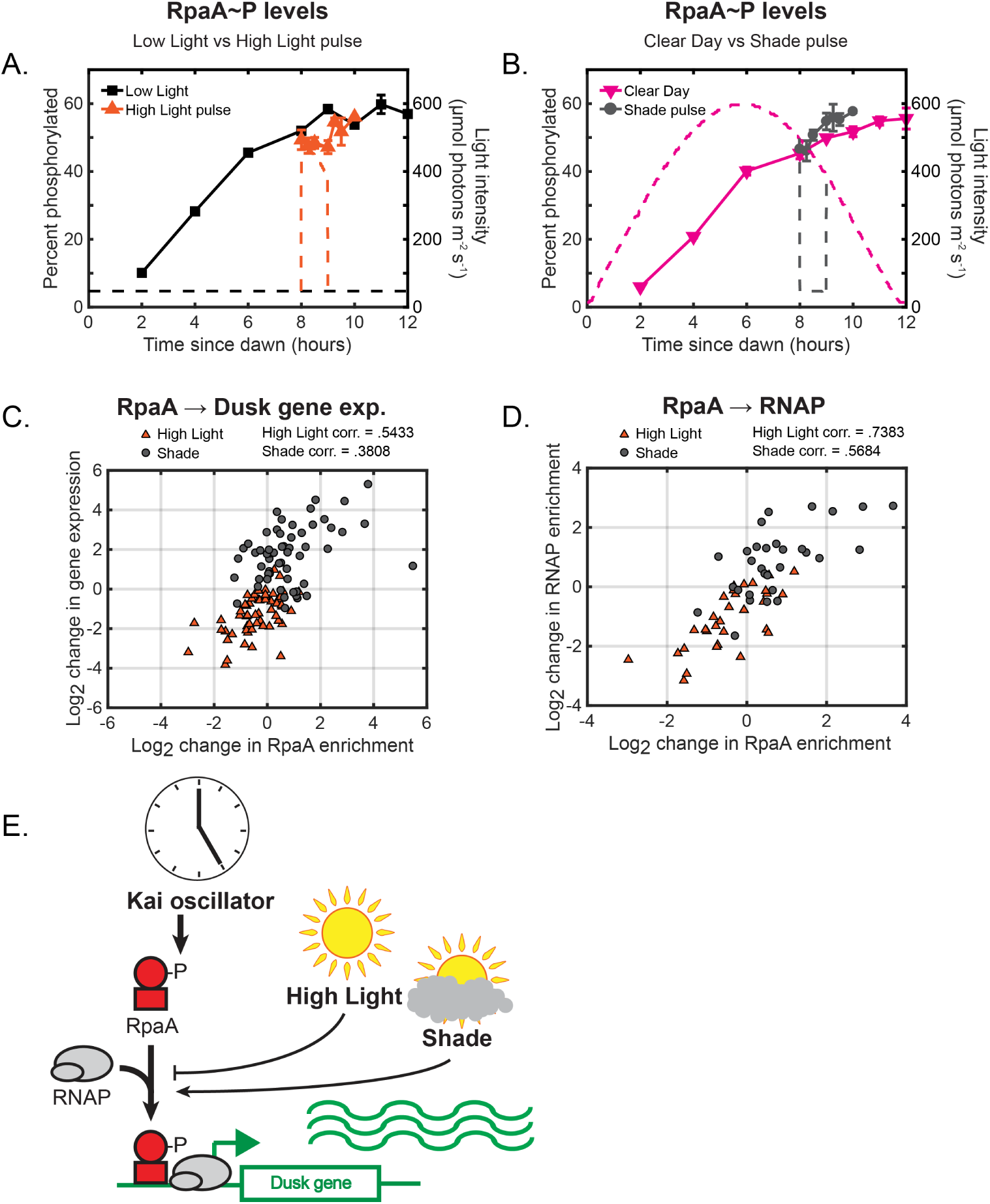
Changes in environmental light intensity regulate RpaA∼P DNA binding activity and RNAP recruitment to control dusk gene expression downstream of clock regulation of RpaA. **(A)** Phosphorylation dynamics of RpaA under Low Light vs High Light pulse. Relative levels of phosphorylated RpaA were measured using Phos-tag Western blotting (left y-axis) in cells grown under Low Light conditions (black squares, see Figure 2A for Setup) or High Light pulse conditions (orange triangles, see Figure 3A for Setup). Each point represents the average of values measured in two independent Western blots, with error bars displaying the range of the measured values. See Materials and Methods, Measurement of RpaA∼P and RpaB∼P levels for more details. Data is available in Figure 4-source data 1. The light profile for each condition is plotted as dashed lines of the same color with values corresponding to the right y-axis. **(B)** Phosphorylation dynamics of RpaA under Clear Day (magenta triangles, see Figure 2A for Setup) vs Shade pulse (gray circles, see Figure 3B for Setup) conditions, measured and plotted as in (A). **(C)** The change in enrichment of RpaA at a given peak upstream of a dusk gene (x-axis) and the corresponding change in expression of the downstream dusk gene (y-axis) from the original condition after 60 minutes in High Light (orange triangles) or Shade (gray circles), plotted for the 56 dusk genes with detectable RpaA peaks in their promoters. The correlation coefficient for the data taken in High Light and Shade conditions is indicated above the plot. See Materials and Methods, ChIP-seq analysis for more details. Data is available in Figure 4-source data 2. **(D)** The change in enrichment of RpaA at a given peak upstream of a dusk gene (x-axis) and the corresponding change in RNAP enrichment upstream of the same gene (y-axis) from the original condition after 60 minutes in High Light (orange triangles) or Shade (gray circles), plotted for the 33 dusk genes with detectable RpaA and RNAP peaks in their promoters. The correlation coefficient for High Light and Shade data is indicated above the plot. See Materials and Methods, ChIP-seq analysis for more details.**(E)** Model of regulation of dusk genes by RpaA under naturally-relevant conditions. The Kai PTO controls levels of RpaA∼P independent of changes in environmental light intensity. Changes in light intensity regulate the recruitment of RpaA∼P with RNAP to dusk genes to control their expression in response to environmental perturbations. (Caption continued on next page.) Figure 4: figure supplement 1 (p. 26). Representative Western blots used to quantify relative levels of RpaA∼P under dynamic light conditions. figure supplement 2 (p. 27). Changes in RpaA enrichment and downstream dusk gene expression after rapid changes in light intensity. figure supplement 3 (p. 27). Changes in RpaA and RNA polymerase enrichment upstream of dusk genes after rapid changes in light intensity. figure supplement 4 (p. 28). Multifactorial behavior of RpaA∼P at select promoters under changes in light intensity. source data 1. Quantification of relative RpaA∼P levels. source data 2. List of RpaA peaks, gene targets, and quantification of enrichment under High Light pulse and Shade pulse conditions.

### D. Phenomenological models suggest simple principles underlying the activation of clusters of light-responsive dusk genes

We find that dusk genes collectively display a small number of responses to changes in environmental light intensity. Using k-means clustering of the gene expression dynamics from our different light profiles (Figures 2,3), as well as from perturbations of RpaA (Figure 2-figure supplement 2 [10]), we identify three major groups of dusk genes (35-80 genes, see Figure 7-source data 1 for full lists) which show distinct and coordinated changes in gene expression over circadian time and in response to changes in light intensity (Figure 7A; Figure 7-figure supplement 1). Under Constant Light conditions, all three clusters are activated by RpaA∼P but display distinct activation dynamics from dawn to dusk (Figure 7-figure supplement 1B) that are mirrored under our Low Light conditions (Figure 7-figure supplement 1A). We named the clusters the Early, Middle, and Late dusk genes based on the order of activation.

The Shade pulse and Sunset in the Clear Day condition have differing effects on the expression of each of the major dusk gene clusters. Early dusk gene expression rapidly increases in response to Shade, but during Sunset plateaus at ∼1/2 of the maximal gene expression reached in Shade (Figure 7A - left plot). Conversely, the Late gene cluster responds most strongly to Sunset in Clear Day conditions but has a mild increase in expression in Shade relative to the Early and Middle dusk genes (Figure 7A - right plot). In contrast, the Middle gene cluster is induced to a similar magnitude by both Shade and Sunset (Figure 7A - middle plot). Shade and Sunset represent similar light changes that occur at different times of day (afternoon and dusk, respectively). As such, the Early and Late dusk gene clusters are differentially induced by a decrease in light intensity depending on the time of day in which it occurs. This circadian effect on the intensity and dynamics of a response to environmental change is a signature of circadian gating [4, 5]. Though circadian gating has been observed (e.g., [35]) and modeled without any knowledge of the transcriptional regulation [36] in plants, it remains unclear what gene regulatory circuits are sufficient to explain such behavior.

At present, there is no mechanistic model to explain the differences in response of these clusters to circadian regulation or changes in sunlight. Given that there are unknown regulators involved in circadian gene expression (Figure 6A,B), and because it is not possible to exhaustively test all possible models of regulation of dusk gene expression, we sought to construct the simplest models that can describe the expression dynamics of these clusters using a phenomenological modeling approach. Such models can be used to highlight regulatory architectures and molecular principles that are sufficient to recapitulate the observed gene expression dynamics, as well as direct further mechanistic studies to reveal the underlying molecular details of gene expression regulation.

Given the clear roles for RpaA∼P and RpaB∼P in activating dusk genes, we asked whether the dynamic expression of the major dusk gene clusters in naturally-relevant light conditions could be described by these variables. We constructed phenomenological models that describe the kinetics of the synthesis and breakdown of an average gene in each of the dusk gene clusters [37] (see Materials and Methods, Mathematical modeling). The rate of synthesis was the sum of a baseline rate of transcription and a maximal adjustable rate of transcription that could be modulated by the activity of one or more regulators. We described the the effects of a regulator such as RpaA∼P or RpaB∼P using a Hill function, whose shape is determined by the Hill coefficient and the coefficient of activation. We determined how well a model could describe the dynamics of a cluster by fitting it to the Clear Day and Shade pulse data and assuming all parameters could vary freely (see Materials and Methods, Mathematical modeling; Table S4).

We began by asking whether the activity of RpaA and/or RpaB can describe the gene expression dynamics of the major dusk clusters in natural light conditions. We first constructed models in which dusk cluster gene expression is solely dependent on RpaA∼P. Activation by RpaA∼P can recapitulate the ordered activation of the dusk gene clusters through differential coefficients of activation for RpaA∼P, but cannot describe the light-responsive expression of these genes (RpaA-only models, Figure 7B-D; Table S5). Further, activation by RpaB∼P alone cannot describe the dusk gene expression patterns of the clusters (RpaB-only models, Figure 7-figure supplement 2; Table S5). However, models in which dusk gene expression is a function of BOTH RpaA∼P and RpaB∼P can recapitulate much of the time-of-day and light intensity dependent expression of the Early and Late clusters and nearly all of the expression dynamics of the Middle clusters (RpaA and RpaB models, Figure 7E-G; Table S5). This suggests that RpaB∼P is a variable which can capture the effects of dynamic light conditions on RpaA∼P activity. The fit parameters for simple joint activation can accommodate indirect activation through the sigma factors and thus do not require direct RpaA/B binding to all genes. Conceptually, our results suggest that transcription factors whose activity track the measured dynamics of both RpaA∼P and RpaB∼P can describe the circadian and light-responsive expression of dusk genes. However, joint activation by RpaA∼P and RpaB∼P predicts that the Early and Late clusters will respond similarly to Shade and Sunset in Clear Day conditions (Figure 7G - left and right plots), and thus cannot capture well the circadian gating of these clusters.

**Figure 5:**
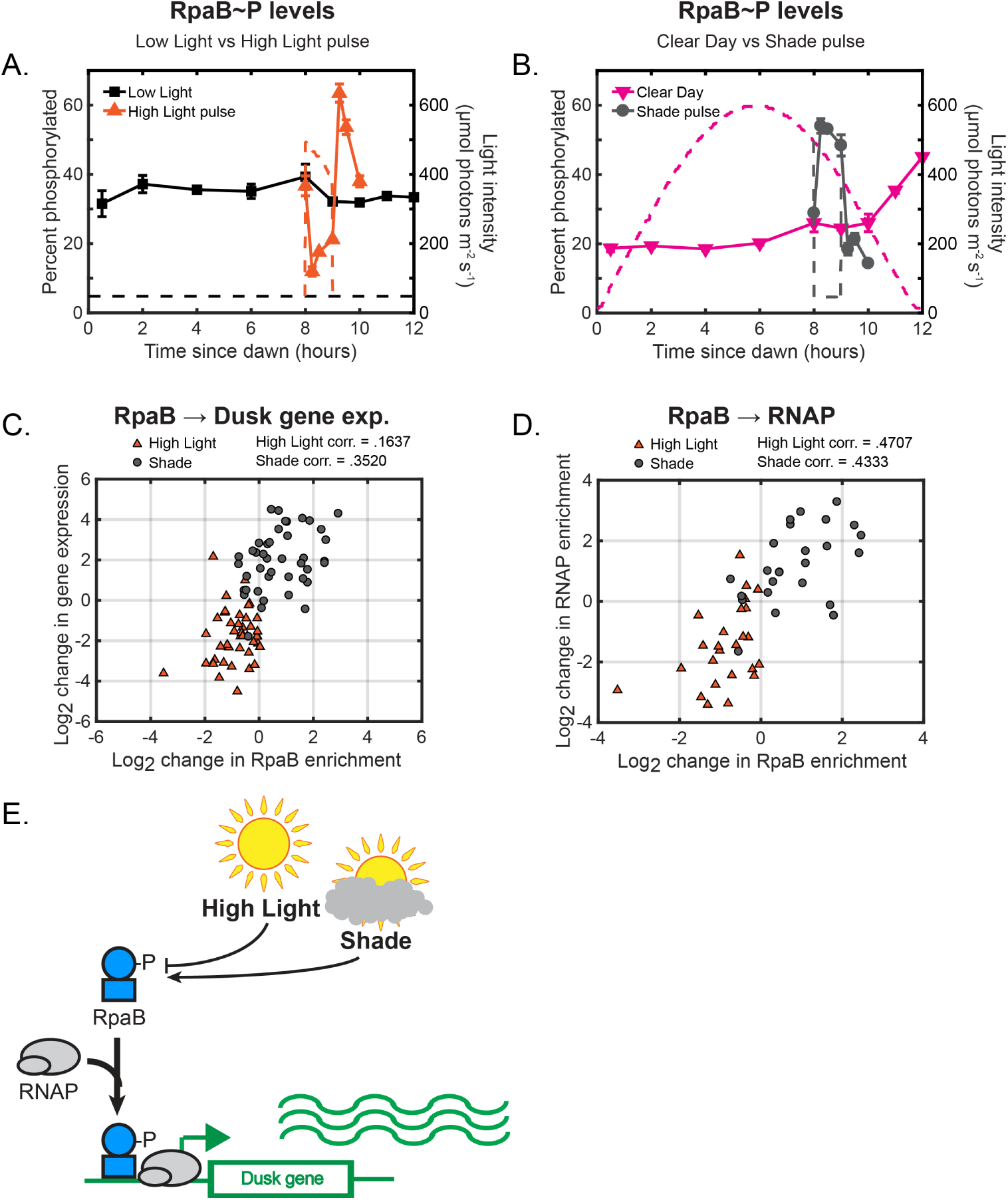
Light-induced changes in RpaB∼P levels modulate RpaB and RNAP binding upstream of dusk genes to directly regulate dusk gene expression in response to light. **(A)** Phosphorylation dynamics of RpaB under Low Light vs High Light pulse. Relative levels of phosphorylated RpaB were measured using Phos-tag Western blotting (left y-axis) in cells grown under Low Light conditions (black squares, see Figure 2A for Setup) or High Light pulse conditions (orange triangles, see Figure 3A for Setup). Each point represents the average of values measured in two independent Western blots, with error bars displaying the range of the measured values. See Materials and Methods, Measurement of RpaA P and RpaB P levels for more details. Data is available in Figure 5-source data 1. The light profile for each condition is plotted as dashed lines of the same color with values corresponding to the right y-axis. **(B)** Phosphorylation dynamics of RpaB under Clear Day (magenta triangles, see Figure 2A for Setup) vs Shade pulse (gray circles, see Figure 3B for Setup) conditions, measured and plotted as in (A). **(C)** The change in enrichment of an RpaB at a given peak upstream of a dusk gene (x-axis) and the corresponding change expression of the downstream dusk gene (y-axis) from the original condition after 60 minutes in High Light (orange triangles) or Shade (gray circles), plotted for the 42 dusk genes with detectable RpaB peaks in their promoters. The correlation coefficient for High Light and Shade data is indicated above the plot. See Materials and Methods, ChIP-seq analysis for more details. Data is available in Figure 5-source data 2. **(D)** The change in enrichment of an RpaB at a given peak upstream of a dusk gene (x-axis) and the corresponding change in RNAP enrichment upstream of the same gene (y-axis) from the original condition after 60 minutes in High Light (orange triangles) or Shade (gray circles), plotted for the 27 dusk genes with detectable RpaB and RNAP peaks in their promoters. he correlation coefficient for High Light and Shade data is indicated above the plot. See Materials and Methods, ChIP-seq analysis for more details. **(E)** Model of regulation of dusk genes by RpaB under naturally-relevant conditions. Changes in light regulate RpaB∼P levels. RpaB∼P binds with RNAP to dusk genes to control their expression in response to environmental perturbations. figure supplement 1 (p. 29). Representative Western blots for quantifing relative levels of RpaB∼P under dynamic light conditions. (Caption continued on next page.) Figure 5: figure supplement 2 (p. 29). Changes in RpaB enrichment and downstream dusk gene expression after rapid changes in light intensity. figure supplement 3 (p. 30). Changes in RpaB and RNA polymerase enrichment upstream of dusk genes after rapid changes in light intensity. source data 1. Quantification of relative RpaB∼P levels. source data 2. List of RpaB peaks, gene targets, and quantification of enrichment under High Light pulse and Shade pulse conditions.

We reasoned that additional regulatory interactions downstream of RpaA and RpaB, or ‘network motifs’ [34], could account for the observed gating of the Early and Late clusters. Thus, we constructed models in which dusk cluster gene expression is positively or negatively dependent on the expression of another cluster alongside activation by RpaA∼P and RpaB∼P (Feedback models, Figure 7H,I; Figure 7-figure supplements 3-5; Table S5). Interestingly, the gating of the Early cluster is recapitulated by a model incorporating an incoherent feedforward loop in which the Late cluster represses Early cluster expression downstream of RpaA∼P and RpaB∼P activation (Figure 7I, left plot; Figure 7-figure supplement 3; Table S5). Further, the gating of the Late cluster is well described by a coherent feedforward loop in which Late cluster expression is dependent on RpaA∼P, RpaB∼P, AND Middle cluster expression levels (Figure 7I, right plot; Figure 7-figure supplement 5; Table S5). Thus, we highlight regulatory schemes downstream of RpaA and RpaB which can generate large time-of-day differences, or circadian gating, in the response to a decrease in light intensity.

Our results highlight that the measured dynamics of RpaA∼P and RpaB∼P can describe the dynamics of large groups of clock-controlled genes after environmental changes and suggest regulatory schemes that can diversify gene expression responses downstream of RpaA and RpaB. The models suggested here offer constraints and testable hypotheses to guide future studies of the molecular mechanisms underlying these responses.

## Discussion

### A. Changes in light adjust circadian gene expression to optimize metabolic changes in response to unexpected and daily changes in light intensity

We show that natural fluctuations in light intensity significantly affect the dynamics of circadian gene expression (Figure 2,3). While previous studies have measured genome-wide gene expression in a single natural light condition [38], here we compare genome-wide circadian gene expression in several physiologically-relevant conditions, including Clear Day, High Light pulse, Shade pulse, and Low Light, to carefully dissect the effects of light on clock output. Natural light changes most greatly affected a large fraction of the dusk genes (Figure 2B and 3C,D), possibly because most of the direct targets of RpaA are dusk genes [10]. We speculate that the opposing trends we observe in dawn gene expression (Figure 2-figure supplement 3 and Figure 3-figure supplement 1) may in part be due to competition for RNAP between the dusk and dawn genes [31, 39] or by growth-rate-dependent mechanisms [32, 33], as this group of genes contains the primary growth genes. A systematic exploration of the effects of light on circadian genes will be necessary to fully elaborate the contributions of light, clock, and growth rate on circadian gene dynamics.

We find that large groups of light-responsive dusk genes are activated by diminished light conditions to different extents depending on the time of day the stimulus is applied, potentially to optimally change metabolism for a given light condition and time of day. The light-responsive dusk genes grouped into three clusters - Early, Middle, and Late - with different activation dynamics during Sunset at the end of the Clear Day versus the Shade pulse in the afternoon (Figure 7A, see Figure 7-source data 1 for full lists of genes in each cluster). Glycogen breakdown genes and the central carbon metabolism genes glyceraldehyde-3-phosphate dehydrogenase and oxalate decarboxylase belong to the Middle dusk genes, which are activated to similar levels by Shade and Sunset. This suggests that cyanobacteria delay the activation of inefficient glycogen breakdown pathways [14] until just before dusk when grown under Clear Day conditions, but can transiently activate these genes in response to Shade to access alternate energy reserves if necessary. Interestingly, genes encoding pyridine nucleotide transhydrogenase, which reversibly converts NADH to the NADPH required for electron transport [40], belong to the Late cluster and are strongly activated only by Sunset and not afternoon Shade. Such a response might allow the delay the adjustment of the relative levels of NADH/NADPH until only when absolutely needed. The cytochrome c oxidase genes belong to the Early cluster, which respond more intensely to Shade than to Sunset (Figure 7A). This enzyme is essential for preventing photodamage in response to rapid changes in light intensity [41]; such changes are not expected to occur during the night, where it serves solely as the terminal electron acceptor for respiration. More generally, the genome-wide gene expression dynamics measured here qualitatively agree with predictions from a whole-cell model of *S. elongatus* that assumed optimization of growth [14]. To resolve how the circadian and light-dependent transcriptional changes effect these metabolic changes, future studies must measure enzyme levels and metabolic fluxes under fluctuating light conditions.

**Figure 6:**
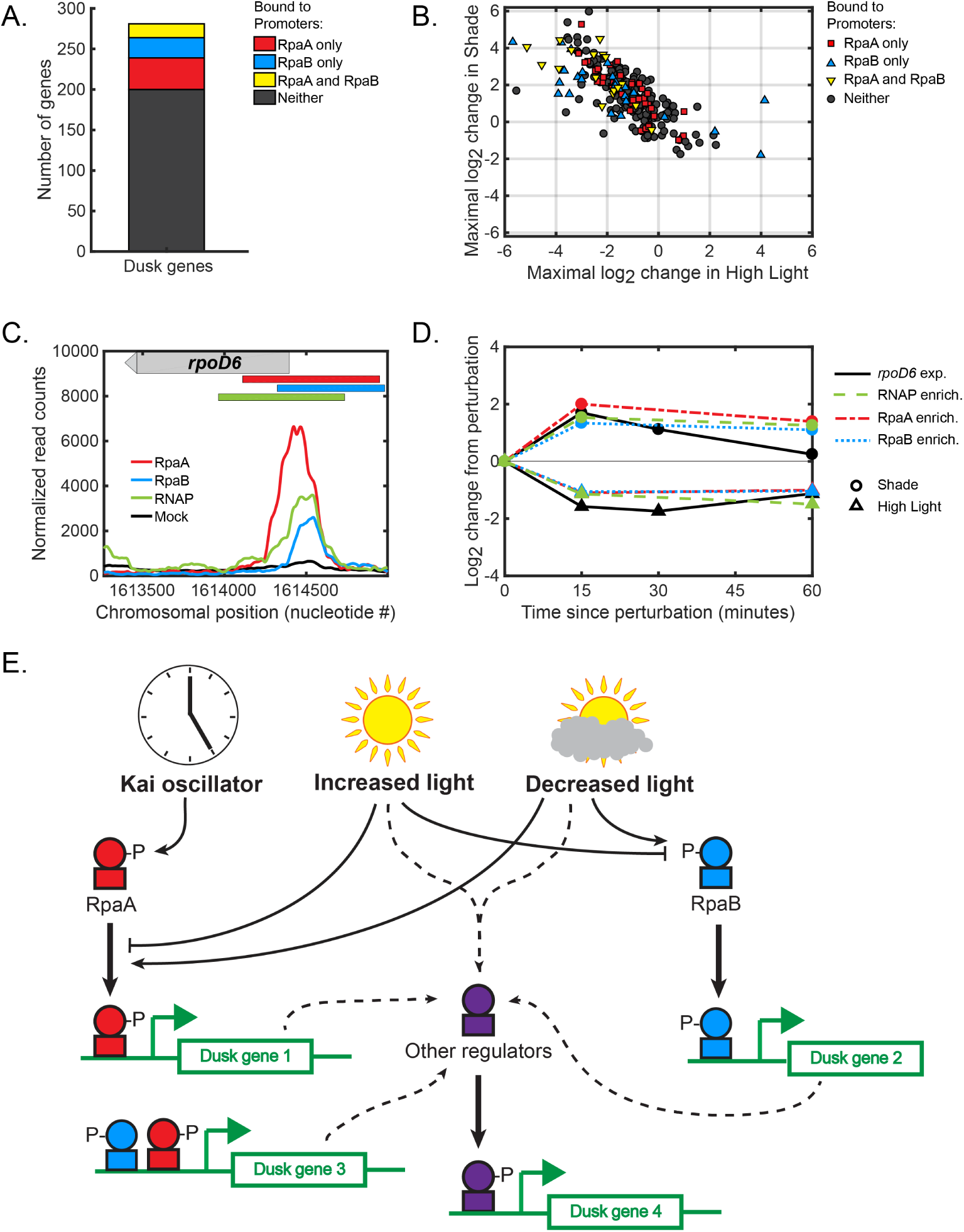
Global regulation of dusk gene expression in response to light changes. **(A)** Number of dusk gene targets of RpaA only (red), RpaB only (blue), RpaA and RpaB (yellow), or neither (black). Target genes of binding sites of RpaA and RpaB were determined using chromatin immunoprecipitation followed by sequencing under several different light conditions (see Materials and Methods, ChIP-seq analysis, for more details. See Figure 2-source data 1 or Figure 3-source data 1 for full lists of RpaA and RpaB peaks associated with dusk genes). **(B)** Light-responsive changes in gene expression of dusk genes. For each dusk gene, we calculated the maximal log_2_ change in expression during the High Light pulse (x-axis) or Shade pulse (y-axis) from 8 hours since dawn in the Low light or Clear day conditions, respectively, using the data from Figure 3. **(C)** Normalized ChIP-seq signal of RpaA (red), RpaB (blue), RNAP (green) and mock IP (black) upstream of the dusk sigma factor gene *rpoD6* at 8 hours since dawn in Low Light. The chromosomal location of the gene is located on the plot with a gray bar with an arrow indicating directionality of the gene. The location of RpaA, RpaB, and RNAP peaks are indicated on top of the plot with red (RpaA), blue (RpaB), and green (RNAP) bars. See Materials and Methods, ChIP-seq analysis for more details. **(D)** Changes in enrichment upstream of *rpoD6* of RpaA (red), RpaB (blue), and RNAP (green) and changes in *rpoD6* gene expression (black) after exposure to the High Light pulse (triangles) or the Shade pulse (circles). See Materials and Methods, ChIP-seq analysis for more details. figure supplement 1 (p. 31). Regulation of dusk sigma factor gene expression by RpaA and RpaB.

**Figure 7:**
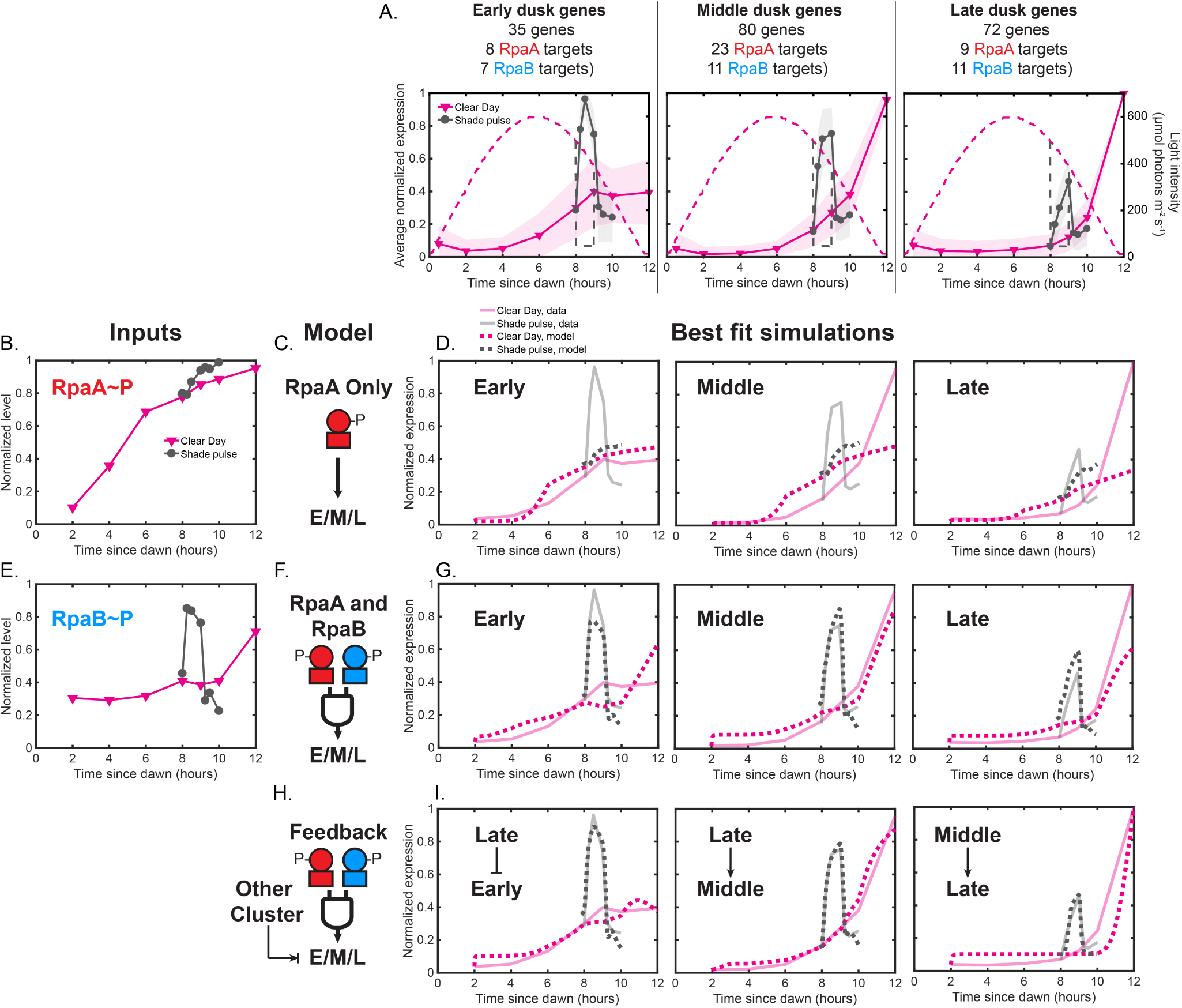
Phenomenological modeling of the activation of clusters of light-responsive dusk genes. **(A)** Average expression profiles of genes within three major clusters of dusk genes under Clear Day (magenta) and Shade pulse (gray) conditions (left y-axis). Dusk genes were grouped using k-means clustering of their normalized expression dynamics in response to the four light conditions of this study and perturbations of RpaA activity in Constant Light conditions (Figure 7-figure supplement 1). Three major clusters emerge - Early (Left plot), Middle (middle plot), and Late (right plot) dusk genes. The number of genes within each cluster, as well as the number of genes with an RpaA or RpaB peak in their promoters (targets) is listed. The expression values of each gene across all four light conditions in this work were normalized to a range of 0 to 1, and the normalized expression values were averaged within each cluster. The shaded region on the plot indicates the standard deviation of the normalized expression values within the cluster. Lists of genes belonging to each cluster and the scaled expression values are available in Figure 7-source data 1. The light intensity profile for each condition is plotted as dashed lines in the same color with values corresponding to the right y-axis. **(B)** Normalized RpaA∼P levels under Clear Day (magenta) and Shade pulse (gray) conditions used as input for mathematical models of dusk gene expression. RpaA∼P levels from all four light conditions were normalized to a range of 0 to 1. **(C)** In the ‘RpaA-only’ models, RpaA∼P activates the expression of the Early (E), Middle (M), or Late (L) cluster. **(D)** Simulations (dotted lines) of best fit RpaA-only models for Clear Day and Shade pulse data (solid lines) for the Early (Left plot), Middle (middle plot), and Late (right plot) dusk genes. The data for each cluster are shown as dotted lines. Data for Clear Day conditions are plotted in magenta, and Shade pulse in gray. **(E)** Normalized RpaB∼P levels under Clear Day (magenta) and Shade pulse (gray) conditions used as input for mathematical models of dusk gene expression. RpaB∼P levels from all four light conditions were normalized to a range of 0 to1. **(F)** In the ‘RpaA and RpaB’ models, RpaA∼P and RpaB∼P jointly activate the expression of the Early, Middle, or Late clusters. **(G)** Simulations of best fit ‘RpaA and RpaB’ models for the Early (Left plot), Middle (middle plot), and Late (right plot) dusk genes, plotted as in (D). (Caption continued on next page.) Figure 7: **(H)** In the ‘Feedback’ models, another cluster activates or represses the expression of the Early, Middle, or Late clusters alongside joint activation by RpaA∼P and RpaB∼P. (I) Simulations (dotted lines) of best fit Feedback models for Clear Day and Shade pulse data (solid lines) for the Early (left plot, Late cluster represses), Middle (middle plot, Late cluster activates), and Late (right plot, Middle cluster activates) dusk genes. The data for each cluster are shown as dotted lines. Data for Clear Day conditions are plotted in magenta, and Shade pulse in gray. figure supplement 1 (p. 32). Average expression profiles of the major dusk gene clusters under various conditions. figure supplement 2 (p. 32). Best fit simulations of ‘RpaB-only’ models in which RpaB∼P solely activates the expression of the dusk gene clusters. figure supplement 3 (p. 33). Models in which either the Middle or Late cluster feeds back to influence Early cluster expression. figure supplement 4 (p. 34). Models in which either the Early or Late cluster feeds back to influence Middle cluster expression. figure supplement 5 (p. 35). Models in which either the Early or Middle cluster feeds back to influence Late cluster expression. source data 1. Lists of genes belonging to the Early, Middle, and Late dusk clusters, and scaled gene expression values.

### B. Mechanistic principles underlying the activation of light-responsive dusk genes

While light does not alter the PTO/TTL circadian circuit, it regulates the activation of dusk genes via RpaA∼P promoter binding (Figure 4) and RpaB promoter binding through its phosphorylation state (Figure 5) at a subset of dusk genes. RpaA binding upstream of its target genes under dynamic light conditions (Figure 4C) correlates with the changes in expression of non-RpaA target genes (Figure 6B). Thus, RpaA∼P may remain the ‘master regulator’ of circadian gene expression whose promoter binding activity is altered by other molecular factors that encode information about the environment, such as RpaB. Previous work suggested that changes in RpaB∼P phosphorylation would alter RpaA∼P levels through competition with the enzymes that control RpaA∼P levels [27]. However, we find that RpaA∼P levels remain constant (Figure 4A,B) under conditions in which RpaB∼P levels change substantially (Figure 5A,B), arguing that RpaB∼P does not influence RpaA∼P levels. RpaB∼P might influence RpaA∼P binding at promoters where both proteins bind (Figure 6-figure supplement 1) as previously suggested [24], and joint control of sigma factors by RpaA and RpaB could feedback to affect RpaA binding at select promoters. Still, the question of how light changes RpaA∼P binding in a promoter-specific way remains unclear.

We define a clear role for the stress-responsive transcription factor RpaB as a transcriptional activator of a large subset of dusk genes (Figure 5E). Further, we demonstrate that decreases in light intensity like a Shade Pulse lead to increases in RpaB∼P levels to allow RpaB to activate the expression of genes, showing that RpaB acts in scenarios beyond its previously appreciated role in High Light stress [25, 42–44]. RpaA∼P and RpaB∼P might cooperate to indirectly regulate the expression of most light-responsive dusk genes by jointly controlling the expression levels of multiple sigma factors (Figure 6-figure supplement 1) [24]. However, our attempts to cleanly perturb RpaB activity to further explore its role as a regulator of dusk genes were unsuccessful, in part because the *rpaB* gene is essential [25]. The role of sigma factors in this network of regulation, while strongly implied, remains ambiguous and attempts to assess this role using genetic deletion of sigma factors yielded inconclusive results. More subtle approaches such as anchors away [45] might allow perturbation experiments that clearly explicate the roles of the sigma factors and RpaB in mediating circadian gene expression.

Although complex molecular mechanisms underlie the light-responsive expression of dusk genes, we demonstrate that phenomenological models effectively describe the differential activation of large groups of dusk genes to afternoon Shade and Sunset. These models suggest that transcription factors with the dynamics of RpaA∼P and RpaB∼P (Figure 7B,E) are sufficient to reproduce much of the activation of the Early, Middle, and Late clusters in response to a Shade pulse in the afternoon or Sunset just before night (Figure 7G). Our models suggest that additional feedback from the other gene clusters may be necessary to achieve the extent of circadian gating observed for the Early and Late clusters (Figure 7I). Our models suggest that interactions between the major dusk clusters can diversify the responses of these clusters to signals from RpaA and RpaB. Regulatory interactions between the sigma factors RpoD6, RpoD5, and SigF2 (Figure 6-figure supplement 1), which belong to the Early, Middle, and Late clusters, respectively, could generate feedback downstream of RpaA and RpaB similar to that in our models (Figure 7-figure supplements 3-5) to generate the diverse responses of the dusk clusters to light conditions. However, feedback could also come from other sources with similar dynamics to the cluster expression levels. Indeed we could not simultaneously fit our models to all four light conditions, likely because of global growth-rate-dependent differences between the Low light and Clear Day conditions. Thus, complete modeling of transcription dynamics of light-dependent dusk genes likely requires explicitly including the effects of metabolism and growth on gene expression [14, 30, 33].

### C. Closing remarks

RpaB and its cognate upstream histidine kinase NblS[46] have been implicated in a variety of stress responses [47–49], which suggests that the mechanisms and regulatory circuits defined here may apply to other environmental changes such as temperature or osmolarity. The requirement of RpaB for mediating the environmental response of circadian genes suggests that the circadian circuit coevolved with RpaB to optimize responses to predictable and unpredictable changes in the environment and motivates the further exploration of the interaction between light and circadian rhythms in *S. elongatus*. Resolution of this interaction and subsequent integration into whole cell models of cyanobacterial growth [30, 50] will help to explain the fitness benefits of the circadian clock [51] and optimize synthetic biology efforts to engineer cyanobacteria to produce useful compounds[52] from the constantly changing sunlight in nature.

## Genomics data

All high throughput data has been submitted to the Gene Expression Omnibus under the title of this manuscript.

## Acknowledgements

JRP and KA thank Phil Shiu, Bin He, Eddie Wang, Andrian Gutu, Chris Chidley, Luca Gerosa, Vadim Patsalo, Rohan Balakrishnan, and Matteo Mori for helpful comments and discussions on the manuscript. JRP thanks Christian Daly, Claire Reardon, Jennifer Couget, and Patrick Dennett of the Harvard Bauer Core Facility for their help with high throughput sequencing and other experiments. KA thanks Al Takeda and Jim MacArthur from the Harvard Electronics Shop for their extensive help in building the controllable lights. JRP and KA were supported by the Howard Hughes Medical Institute through EKO.

## Materials and Methods

### Cyanobacterial strains

Most experiments were conducted in a pure wildtype background of *Synechococcus elongatus* PCC7942 (ATCC catalog number 33912). For RNAP ChIP experiments, we used a strain in which the *β′* subunit of RNA polymerase (*Synpcc 7942_1524*) was C-terminally tagged with a 3x FLAG epitope (a gift from Ania Puszynska). To make this strain, wildtype *S. elongatus* was transformed with a plasmid encoding the *Synpcc 7942_1524* gene with sequence encoding a 3X GS linker and a 3X FLAG epitope inserted before the stop codon, targeted to insert at the native locus of the gene. A downstream kanamycin resistance cassette was used for selection. This plasmid is available through Addgene with the ID 102337. Two different clones of this strain, EOC398 and EOC399, were confirmed by sequencing colony PCR fragments that amplified the modified regions of the gene, and the presence of the tagged subunit was confirmed with Western blotting.

### Construction of light apparatus

To grow the cyanobacteria in different light profiles, we constructed an apparatus to control the intensity of four high powered LED arrays (parts list in Table S1, p. 21). ‘Warm white’ LED arrays (∼1 in. x 1 in., Bridgelux) were chosen because of maximal overlap with the phycobilisome absorption spectrum. An LED array was mounted on a heatsink (Nuventix) and powered by a Flexblock LED driver (LEDdynamics) wired in the ‘boost only’ configuration (Table S2, p. 21). The intensity of the LEDs was controlled by varying the voltage input into the DIM line of the Flexblock between 0 and 10 V. We used a digital potentiometer (AD7376, Analog Devices) as a controllable 10 V source. The voltage output of the digipot was controlled via serial peripheral interface with an Arduino Uno board (Arduino) (see Table S3, 23). Each LED array was controlled separately, and a single array was sufficient to grow a single 750 mL culture of *S. elongatus*. All wires carrying substantial currents from the main power supply to the LED arrays were rated 18 AWG, and all other wires were rated 22 AWG. The relatively low voltage of the main power supply (18 V) is essential for being able to turn off the LED arrays completely.

### Calibrating light conditions

A single LED was mounted to shine perpendicular to the ground and isolated from other light sources. A single 750 mL cyanobacterial culture in a 150 cm^2^ BD Falcon Tissue culture flask (Fisher Scientific) was placed beneath the LED, tilted such that the broad face of the culture was almost perpendicular to the incoming light. Each LED was calibrated by passing a known voltage input to the LEDs and recording the intensity of the light in *µ*mol photons m^−^2 s^−^1 at the position of the surface of the culture directly beneath the LED using a LI-COR LI-250A light meter equipped a quantum sensor. To access a greater dynamic range of light intensity values, we calibrated the lights to give light intensity values at either of two distances from the light source — raised towards the lights to access higher light intensities, or lowered away from the lights to access lower light intensities.

To define the Clear Day conditions, we used light intensity values measured by the Ground-based Atmospheric Monitoring Instrument Suite, Rooftop Instrument Group on March 23rd, 2013 (Figure 1B, dark blue line, [6]). We used this light intensity profile to define the rate of change of light intensity in our Clear Day condition, with a maximal light intensity of 600 *µ*mol photons m^−2^ s^−1^. This intensity is consistent with measurements of light intensity in aquatic environments [38], while also offering an order of magnitude difference in intensity compared to the Low Light condition, which was a constant 50 *µ*mol photons m^−2^ s^−1^. The Shade pulse condition was defined by dividing the intensity value of our Clear Day profile by 10 fold between 8 and 9 hours after dawn. The High Light pulse was defined as the intensity of the Clear Day condition between 8 and 9 hours after dawn. Low Light cultures were grown continuously at 50 *µ*mol photons m^−2^ s^−1^. We generated the dynamic changes in light intensity of our conditions by changing the intensity of the LED every three minutes by passing the calibrated voltage value corresponding to the appropriate light intensity of our defined profile. After the 12-h light profile, the LEDs were turned off for 12 hours during the dark period. Cultures were grown semi-turbidostatically (OD_750_ maintained at 0.3) with periodic dilution in BG-11M media supplemented with 10 mM HEPES pH 8.0 at 30 °C,continuously bubbled with 1% CO_2_ in air, and shaken at 25 rpm in an enclosure impermeable to room lighting. Cells were not grown with antibiotics during the course of the experiment.

### Purification of anti-RpaA and anti-RpaB antibodies

Recombinant RpaA was purified as previously described [18]. To purify recombinant RpaB, we cloned the *rpaB* gene into the pET48-b+ plasmid (Novagen) and overexpressed Trx-His-tagged RpaB in Novagen Tuner (DE3) competent cells carrying this plasmid by adding 300 *µ*M IPTG to mid-log phase cultures. RpaB was purified from cell lysate using Ni-NTA chromatography as described previously [19]. The Trx-His tag was cleaved from RpaB and removed using a subsequent Ni-NTA step as described [19]. Purified, cleaved RpaB was dialyzed into a buffer containing 20 mM HEPES-KOH, pH 8.0, 150 mM KCl, 10% w/v glycerol, and 1 mM DTT. Protein concentration was measured with the Pierce BCA assay, and aliquots were flash frozen and stored at −80°C.

Anti-RpaB serum was generated by immunization of two rabbits with purified RpaB by Cocalico Biologicals (Reamstown, PA). RpaA- and RpaB-conjugated Affigel 10/15 resin (Bio-Rad) was prepared following manufacturer’s instructions as described previously [19]. Anti-RpaB serum was first passed over an RpaA-conjugated resin and the flowthrough collected to subtract cross-reacting antibodies. Anti-RpaB antibodies were then purified from the flowthrough using an RpaB-conjugated resin as described previously [19]. The same process was repeated to purify anti-RpaA antibodies using rabbit serum described previously [10], passing the serum over an RpaB-conjugated resin and purifying with an RpaA-conjugated resin. No cross reactivity of the purified anti-RpaA and anti-RpaB antibodies for the opposite regulator was detected via ELISA assay.

### Measurement of RpaA∼P and RpaB∼P levels

Ten mL of cyanobacterial culture with OD_750_ = 0.3 were collected on cellulose acetate filters and flash frozen prior to storage at −80 ^°^C. Cell lysates for Western blotting were prepared from the collected cells as described previously [10]. Equal amounts of cell lysate (10-15 *µ*g) were resolved on Phos-tag acrylamide gels and transferred to nitrocellulose membranes as described previously [19]. Membranes were probed with 1/5000 dilution of purified anti-RpaA and anti-RpaB antibody. RpaA blots were then incubated with goat anti-rabbit HRPconjugated secondary antibody and developed using the Pierce Femto chemiluminescence kit. The exposed blots were imaged with an Alpha Innotech Imaging station. RpaB blots were incubated with Goat anti-Rabbit Westerndot 585 antibody and imaged with a Typhoon Imager. The intensities of the bands corresponding to unphosphorylated and phosphorylated RpaA/B were quantified using Imagequant software (GE Healthcare Life Sciences) using rubber band background subtraction. The percent of RpaA (or RpaB) phosphorylated was quantified as the intensity of the RpaA∼P band divided by the sum of the intensities of the RpaA and RpaA∼P bands, multiplied by 100. Values reported in Figures 4A,B and 5A,B represent the average of two separate measurements from replicate Western blots, with error bars displaying the range of the measured values (See Figure 4-source data 1, and Figure 5-source data 1 for raw data from the replicate experiments). The trends seen were reproducibly observed between separate biological replicates of the light condition time courses.

### RNA sequencing

Twenty-five mL of cyanobacterial culture with OD_750_ = 0.3 were collected on cellulose acetate filters and flash frozen prior to storage at −80 ^°^C. Cells were resuspended in RNAprotect Bacteria reagent (Qiagen), and 1/3 of the cells were resuspended in a buffer containing 15 mg/mL lysozyme, 10 mM Tris-Cl, 1 mM EDTA pH 8, and 50 mM NaCl and incubated for 10 minutes. RNA was purified from the lysed cells using the Qiagen RNeasy Mini Kit. Ribosomal RNA was depleted from1.25 *µ*g of purified RNA using the Ribo-Zero bacteria rRNA removal kit (Illumina). Strand-specific RNAseq libraries were prepared from the depleted RNA using the Truseq Stranded mRNA Sample prep kit (Illumina) and sequenced on an Illumina HiSeq 2500 machine by the Bauer Core Facility at the Harvard FAS Center for Systems Biology. Sequencing reads were aligned to the *S. elongatus* genome as described previously [10], with samples averaging 8 million aligned reads. We quantified expression of a gene by counting the number of aligned sequencing reads corresponding to the appropriate strand between the start and stop of each gene, and normalized these values between all samples from the light conditions in this work using median normalization, followed by dividing the median normalized read count value by the length of the open reading frame of the gene, as described previously [10, 53]. The time course and RNA sequencing was repeated twice for two biological replicates (data available in Figure 2-source data 1 and Figure 3-source data 1). The data plotted in this work are from replicate 2, and the trends observed are reproduced in both biological replicates.

### Definition of circadian genes

We defined a subset of previously identified circadian genes on which to focus our analysis. We began with a list of 856 previously described reproducibly circadian genes [8, 10]. We next required that these genes have a Cosiner amplitude [54] of greater that 0.15 under Constant Light conditions [8]. We also required that the gene display expression of at least 1 read per nucleotide in at least one time point of the RNA sequencing experiments in this study. These filters produce a list of 450 high confidence circadian genes.

We noted that genes classified as dawn (class 2) and dusk (class 1) genes under Constant Light conditions [8] showed maximal expression at a different time of day under our Low Light conditions, while the relative ordering of genes by Cosiner phase [54] from Constant Light conditions [8]was preserved. As such, we redefined dawn genes as those genes with a phase of 40° to 189° under Constant Light conditions [8], and dusk genes as those with a phase of 190° to 360° and 0° to 39°, as determined by the Cosiner algorithm [54]. These definitions produce a list of 169 high confidence dawn genes, and 281 high confidence dusk genes. The expression of our redefined circadian genes under Constant Light conditions is plotted in Figure 2-figure supplement 2. The list of high confidence circadian genes and high confidence class assignments is available in Figure 2-source data 1 and Figure 3-source data 1.

### ChIP sequencing

One hundred and twenty mL of OD_750_ 0.3 cyanobacterial culture were removed and crosslinked with 1% formaldehyde at 30 °C for 5 minutes in front of a light source. Crosslinking was quenched with 125 mM glycine. Crosslinked cells were washed twice with phosphate buffered saline, pelleted, and flash frozen prior to storage at −80 ^°^C.

Pellets were resuspended in 1 mL of BG-11M supplemented with 500 mM L-proline and 1 mg/mL lysozyme and incubated at 30 ^°^C for 1 hour to digest the cell wall. Cells were collected and resuspended in a Lysis buffer (50 mM HEPES pH 7.5, 140 mM NaCl, 1 mM EDTA, 1% Triton X-100, 0.1% sodium deoxycholate, and 1x Roche Complete EDTA-free Protease Inhibitor Cocktail) prior to shearing in a Covaris E220 Adaptive Focus System (Peak Incident Power = 175; Duty Factor = 10%; Cycles per burst = 200; Time = 160 s). The lysates were cleared via centrifugation, and concentration was determined via the Pierce BCA Assay.

For a given pulldown, 800 *µ*g of lysate was incubated overnight at 4 ^°^C in 500 *µ*L of lysis buffer with 8 *µ*g of anti-RpaA, anti-RpaB, or FLAG M2 mouse monoclonal antibody (Sigma-Aldrich) for RNAP pulldowns. A mock pulldown was carried out in which equal amounts of lysate from every time point of the time course (Shade 0, 15, 60 minutes, High Light 0, 15, 60 minutes) in a total of 800 *µ*g was incubated with 8 *µ*g of rabbit Igg. Next, 35 *µ*L of Dynabeads protein G (Thermo Fischer Scientific) equilibrated in lysis buffer were added and the sample was incubated with mixing for 2 hours at 4 ^°^C.The beads were washed and DNA was eluted and purified as described previously [10].

Sequencing libraries were prepared from the purified ChIP DNA using the NEBNext Ultra II DNA Library Prep Kit (New England Laboratory, Ipswich, MA). Libraries were sequenced on an Illumina HiSeq2500 instrumentby the by the Bauer Core Facility at the Harvard FAS Center for Systems Biology. We created sequencing libraries of ChIP experiments from two separate biological repeats of the time course experiment. Reads were aligned to the *S. elongatus* genome as described previously [10], resulting in an average of 3 million aligned reads for replicate 1, and 5 million aligned reads for replicate 2.

### ChIP-seq analysis

The aligned read data per genomic position was smoothed with a Gaussian filter (window size = 400 base pairs, standard deviation = 50). Each data set was normalized to the Mock ChIP-seq experiment and peaks which were significantly enriched above the Mock were identified in each data set using a previously described[10] custom-coded form of the Peak-seq algorithm [55]. Within each replicate time course for a given protein, we compiled a list of peaks which were enriched at least 3.5 fold over the Mock experiment at the position of highest ChIP signal. Finally, we required that a peak be detected in both replicates for it to be considered. This analysis generated 114 RpaA peaks, 218 RpaB peaks, and 451 RNAP peaks. To calculate enrichment for a peak, we determined the ChIP signal at a given time point at the genomic position of the highest ChIP signal detected for that peak and divided this by the value of the Mock experiment at that position. The data plotted in this manuscript are from replicate 2, but all trends hold in replicate 1. We assigned a gene as a target of a peak if: i) the start codon of the gene was within 500 bp of the position of maximal ChIP signal within a peak; ii) the peak resided upstream of the gene; iii) The gene was the closest gene to that peak on the same strand. Lists of RNAP, RpaA, and RpaB peaks and gene targets are found in Figure 2-source data 2, Figure 4-source data 2, and Figure 5-source data 2, respectively.

For Figure 2G, 3C, and 4C, we identified all RNAP, RpaA, or RpaB peaks with dusk gene targets based on the above criteria, respectively. 82 dusk genes are targets of RNAP peaks, 56 dusk genes were targets of RpaA peaks, and 42 dusk genes are targets of RpaB peaks. Then, for each peak - dusk gene pair, we calculated the change in gene expression of the dusk gene after 60 minutes, and the change in ChIP enrichment of the upstream peak over the mock pulldown (described above) after 60 minutes in High light, each compared to their respective values at Low light at 8 hours since dawn. We plotted these data on the x- and y-axes, respectively, with orange triangles. We repeated this process, comparing gene expression and ChIP enrichment values after 60 minutes in Shade compared to 8 hours since dawn in Clear Day conditions, and plotted the data as gray circles. We calculated the correlation coefficient between the change in gene expression and the change in ChIP enrichment for all peak-gene pairs of the relevant factor in the High Light pulse, and then calculated the same correlation in Shade pulse conditions separately. We calculated the correlation coefficients comparing changes after 15 minutes in either the High Light or Shade pulse conditions, and list these values in the legends of Figure 3-figure supplement 2, Figure 4-figure supplement 2, and Figure 5-figure supplement 2. The data used for these plots for RNAP, RpaA, and RpaB are available in Figure 3-source data 2, Figure 4-source data 2, and Figure 5-source data 2, respectively. We plot data from replicate 2, and the trends are reproduced in replicate 1.

For Figure 3-figure supplement 2, Figure 4-figure supplement 2, and Figure 5-figure supplement 2, we took the lists of RNAP/RpaA/RpaB peaks with dusk gene targets from above. For each peak - gene pair, we calculated the log_2_ fold change in ChIP enrichment of the peak and the change in expression of the downstream gene in 15 or 60 minutes in the High Light pulse compared to the value at 8 hours since dawn in Low Light conditions. We repeated these calculations for each peak-gene pair in 15 or 60 minutes in Shade pulse compared to 8 hours since dawn in Clear Day conditions. We used hierarchical clustering on the collective ChIP and gene expression data from both conditions to determine the plotting order of the peak-gene pairs in the heat maps, and then plotted the log_2_ change in ChIP enrichment and dusk target gene expression in the two conditions in separate heat maps. The change in enrichment of a peak and the change in expression of its target dusk gene are aligned horizontally in their respective heat maps. The leftmost column of each heat map is white, because this column compares the time 0 data to itself and thus has a log_2_ value of 0. One RpaA peak resides upstream of two dusk genes, and two RpaB peaks reside upstream of two dusk genes each, and thus the listed number of RpaA and RpaB peaks is smaller than the number of RpaA and RpaB target dusk genes. The data used for these plots for RNAP, RpaA, and RpaB are available in Figure 3-source data 2, Figure 4-source data 2, and Figure 5-source data 2, respectively. We plot data from replicate 2, and the trends are reproduced in replicate 1.

For Figure 4D and 5D we identified all dusk genes that were targets of both RpaA and RNAP (for figure 5D) or both RpaB and RNAP (for Figure 5D). 33 dusk genes are targets of both RpaA and RNAP peaks, and 27 dusk genes are targets of both RpaB and RNAP. Then, for each pair of RpaA/B - RNAP peaks, we calculated the change in ChIP enrichment of the RpaA/B peak after 60 minutes, and the change in ChIP enrichment of the RNAP peak upstream of the same dusk gene over the mock pulldown (described above) after 60 minutes in High light, each compared to their respective values at Low light at 8 hours since dawn. We plotted these data on the x- and y-axes, respectively, with orange triangles. We repeated this process, comparing RpaA/B ChIP enrichment and RNAP ChIP enrichment values after 60 minutes in Shade compared to 8 hours since dawn in Clear Day conditions, and plotted the data as gray circles. We calculated the correlation coefficient between the change in RpaA/B ChIP enrichment and the change in RNAP ChIP enrichment for all RpaA/B - RNAP peak pairs of the relevant factor in the High Light pulse, and then calculated the same correlation in Shade pulse conditions separately. We calculated the correlation coefficients comparing changes after 15 minutes in either the High Light or Shade pulse conditions, and list these values in the legends of Figure 4-figure supplement 3, and Figure 5-figure supplement 3. The RNAP, RpaA, and RpaB peaks associated with each dusk gene are listed in Figure 2-source data 1 and Figure 3-source data 1, and the enrichment values for these peaks are listed in Figure 3-source data 2, Figure 4-source data 2, and Figure 5-source data 2, respectively. The data plotted here are from replicate 2, and the trends are reproduced in replicate 1.

For Figure 4-figure supplement 3 and Figure 5-figure supplement 3, we took the lists of RpaA/RpaB - RNAP peaks pairs upstream of the same dusk gene from above. For each RpaA/B - RNAP peak, we calculated the log_2_ fold change in ChIP enrichment of the RpaA/B peak and the change in ChIP enrichment of the RNAP peak upstream of the same dusk gene in 15 or 60 minutes in the High Light pulse compared to the value at 8 hours since dawn in Low Light conditions. We repeated these calculations for each peak-gene pair in 15 or 60 minutes in Shade pulse compared to 8 hours since dawn in Clear Day conditions. We used hierarchical clustering on the collective RpaA/B and RNAP ChIP data from both conditions to determine the plotting order of the RpaA/RpaB - RNAP peak pairs in the heat maps, and then plotted the log_2_ change in RpaA/B ChIP enrichment and RNAP ChIP enrichment in the two conditions in separate heat maps. The change in enrichment of an RpaA/B peak and the change in enrichment of the RNAP peak upstream of the same dusk gene are aligned horizontally in their respective heat maps. The leftmost column of each heat map is white, because this column compares the time 0 data to itself and thus has a log_2_ value of 0. The RNAP, RpaA, and RpaB peaks associated with each dusk gene are listed in Figure 2-source data 1 and Figure 3-source data 1, and the enrichment values for these peaks are listed in Figure 3-source data 2, Figure 4-source data 2, and Figure 5-source data 2, respectively. The data plotted here are from replicate 2, and the trends are reproduced in replicate 1.

For Figure 4-figure supplement 4D-F, Figure 6D, and Figure 6-figure supplement 1D-F, we identified all RpaA, RpaB, and RNAP peaks that targeted the specified gene, as described above. Then, we calculated the log_2_ change in RpaA (red dashed line), RpaB (blue dotted line), RNAP (green dashed line) ChIP enrichment or expression of the downstream gene (black solid lines) in the High Light pulse compared to 8 hours since dawn in the Low Light condition, and plotted these values with downward triangles. We repeated these calculations, comparing enrichment and gene expression in the Shade pulse to the data at 8 hours since dawn in the Clear Day condition, and plotted these values with circles. The RNAP, RpaA, and RpaB peaks associated with each dusk gene are listed in Figure 2-source data 1 and Figure 3-source data 1, and the enrichment values for these peaks are listed in Figure 3-source data 2, Figure 4-source data 2, and Figure 5-source data 2, respectively. The data plotted here are from replicate 2, and the trends are reproduced in replicate 1.

### K-means clustering

We calculated normalized expression values of high confidence dusk genes under our dynamic light conditions, as well as in previously described RpaA perturbations in Constant Light [10]. We separately normalized the data from set of dynamic light conditions (Low Light, Clear Day, High Light pulse, Shade pulse) and the Constant Light data (Wildtype, OX-D53E cells — *rpaA-*, *kaiBC-*, *Ptrc::rpaA(D53E)* — without inducer, OX-D53E with inducer) using z-score normalization, and used this data to separate the dusk genes into 8 groups with k-means clustering using Pearson correlation as the distance metric. We focused our analysis on the three largest clusters which accounted for most of the dusk genes (187/281 genes). The lists of genes belonging the three major clusters are found in Figure 7-source data 1.

### Mathematical modeling

We observed very regular and systematic changes in the expression of large clusters of dusk genes in natural light conditions (Figures 2, 3, and 7A) that correlated with RpaA/B recruitment of RNAP (Figures 4-6). Thus, our goal was to determine whether simple phenomenological models similar to that inspired by Alon[34] could reproduce these observations and offer some intuition into how they might arise. Such a model does not take into account the specific molecular mechanisms underlying RpaA/B regulation explored in the Results. Additionally, while most of the dusk genes underwent systematic changes, a small group of ∼20 genes including *kaiBC* was relatively insensitive to changes in light intensity (Figure 4-figure supplement 4), and we do not model those genes’ expression dynamics.

Our model treats the activation or repression of the expression of a dusk gene cluster by RpaA∼P, RpaB∼P, or another cluster using effective Hill kinetics. We coarse-grained each of the three groups of circadian dusk genes (the Early, Middle, and Late clusters in Figure 7A) to a single effective gene with the average dynamics of the group (Figure 7A, triangles). We modeled the dynamics of a gene cluster X using a simple kinetic model of an AND gate at a promoter [37],

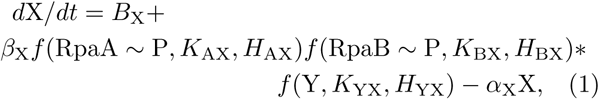

where *B*_X_ is the basal transcription rate; *f* is a function of the interaction of X with RpaA∼P, RpaA∼P, or another cluster Y; *β*_X_ is the max transcription rate; and *α*_X_ is the decay/dilution rate. Activating interactions were treated using a simple Hill function,

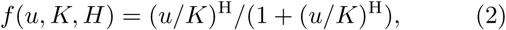

where *u* is the concentration of the active transcription factor, H is the Hill coefficient of interaction, and *K* is the coefficient of activation. Bacteria can easily tune the interactions between proteins and between transcription factors and promoters to adjust H and *K* for different clusters [56]. RpaA∼P and RpaB∼P were treated as activators, consistent with the results from Figures 4-6. Repressive interactions between clusters were treated using

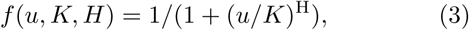

where *K* is now the coefficient of repression. In equation 1, RpaA∼P, RpaB∼P, and Y were measured experimentally; the remainder of the parameters were left free.

We determined the sufficiency of a model to describe the data by fitting the parameters using the range of values shown in Table S4. Time propagation of the differential equation 1 was performed using the *ode45* solver in MATLAB, with X(*t* = 0) set as the observed expression level at the beginning of the simulated time period. Model fitting was performed in MATLAB using the nonlinear least squares solver *lsqnonlin*.

The Akaike Information Criterion (AIC) and the Chi-squared test are typically used to quantify whether a model with more parameters fits the data better than another with fewer parameters simply because it is more complex. However, both approaches are for statistical models in which little to no information is used to construct the model and are not strictly applicable to the model constructed here, which is based on our understanding of transcription. If we do use AIC to compare the models, the feedback models are predicted to be most probable.

In our model, H and *K* are effective constants that represent the overall ability of RpaA∼P, RpaB∼P, or another gene cluster Y to affect gene expression. These constants include potential indirect activation through the sigma factors, which is may be why joint activation by RpaA∼P and RpaB∼P describe the dynamics of the Middle cluster reasonably well. However, circadian gating of the Early and Late dusk genes requires further interactions that cannot be described by Hill functions of measured RpaA∼P and RpaB∼P levels. Clearly there may be more complex networks at play that those we have considered here, and much more needs to be done to fully model gene expression in *S. elongatus*. Here we have constructed a first model to suggest simple principles underlying the interaction of circadian and light regulation of dusk genes and offer directions for further exploration.

**Figure 2-figure supplement 1:**
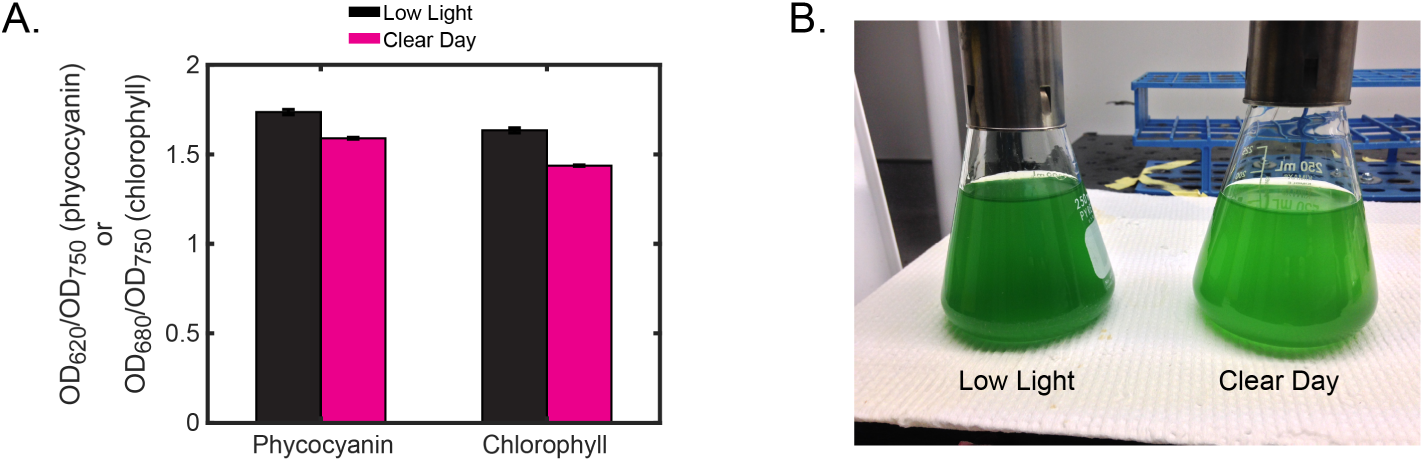
Pigment levels of cyanobacteria grown under Low Light or Clear Day conditions reveal adjustments in the photosynthetic apparatus to optimize growth in different light conditions. **(A)** Estimation of phycocyanin and chlorophyll levels in cells grown under Low Light (black) or Clear Day (magenta) conditions for two days, measured at midday of the third light period. Phycocyanin and chlorophyll levels were estimated by measuring optical density of the culture at 620 nm or 680 nm, respectively, and normalizing to optical density at 750 nm to account for differences in cell density. Error bars show the standard deviation of three independent measurements. Cells grown under Clear Day conditions show lower levels of both phycocyanin and chlorophyll. **(B)** Image of cells harvested from cultures at midday on the third day (OD_750_ = 0.3) following two 12-h light/12-h dark cycles in either Low Light (left) and Clear Day (right) conditions (see Figure 2A). The more yellow-green color of the cells from Clear Day conditions compared to the Low Light grown cells, which are blue-green, indicates diminished levels of phycocyanin. Cells grown in Clear Day divided roughly twice as fast at midday compared to Low Light cells (∼6 hour doubling in Clear Day compared to ∼12 hour doubling time in Low Light). The growth rate of cells grown in Clear Day conditions varied throughout the day, showing slowest growth just prior to dusk (∼10 hour doubling time). Growth rate was estimated by calculating the change in the ln(OD_750_) of the culture before and after a 1 hour time period. Taken together, the growth and pigment differences suggest that the cells lowered the levels of photosynthetic components to cope with the increased photon flux of Clear Day, while using these additional resources for faster growth.

**Table S1:** Parts for controllable light source. The table includes the parts chosen for their specific properties. The remaining parts, such as wires, heat shrink tubing, thermal paste for mounting the LEDs on the heat sinks, proto-boards, and housing are quite general and specific brands are unnecessary.

| Part name | Digikey part number | Current price (\$) | Quantity |
| --- | --- | --- | --- |
| PWR SUP MEDICAL 18V 8.3A 150W | EPS439-ND | 73.71 | 1 |
| CONN RCPT 8CONT DIN SLD PNL MNT | SC2007-ND | 5.64 | 1 |
| LEDDynamics Flexblock BUCK BOOST 48V, 700 mA | 788-1038-ND | 19.99 | 4 |
| AD7376 digital potentiometer | AD7376ARWZ10-ND | 8.66 | 4 |
| AC to DC power supply, 10VDC, 275 mA | 993-1233-ND | 4.68 | 2 |
| BXRA-30E1200-B-03, Bridgelux, Warm white, LED | Not sold at Digikey.<br>Need to order from:<br>AMBIT ELECTRONICS, INC. | 10.47 | 4 |
| Aavid thermalloy Spotlight 47W heat sink | 1061-1092-ND | 9.50 | 4 |
| Arduino Uno Board Rev3 | 1050-1024-ND | 21.49 | 1 |

**Table S2:**
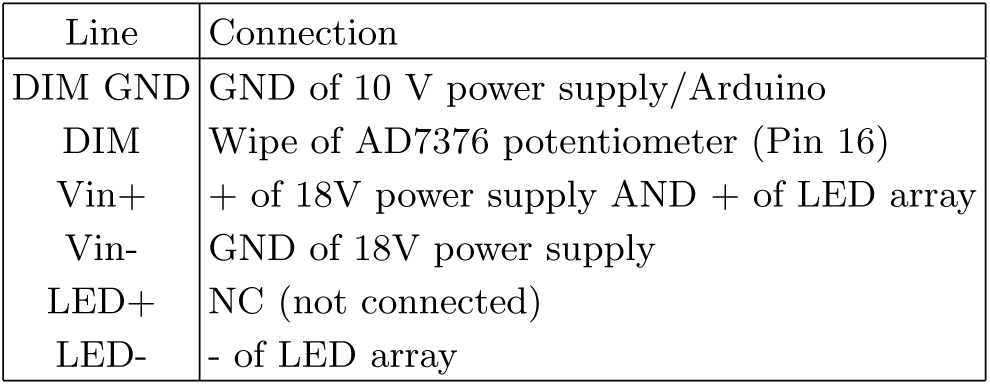
Wiring the FlexBlock LED driver. The FlexBlock LED driver needs to be connected in a ‘boost only’ configuration (see spec sheet for more details), with connections as shown.

**Figure 2-figure supplement 2:**
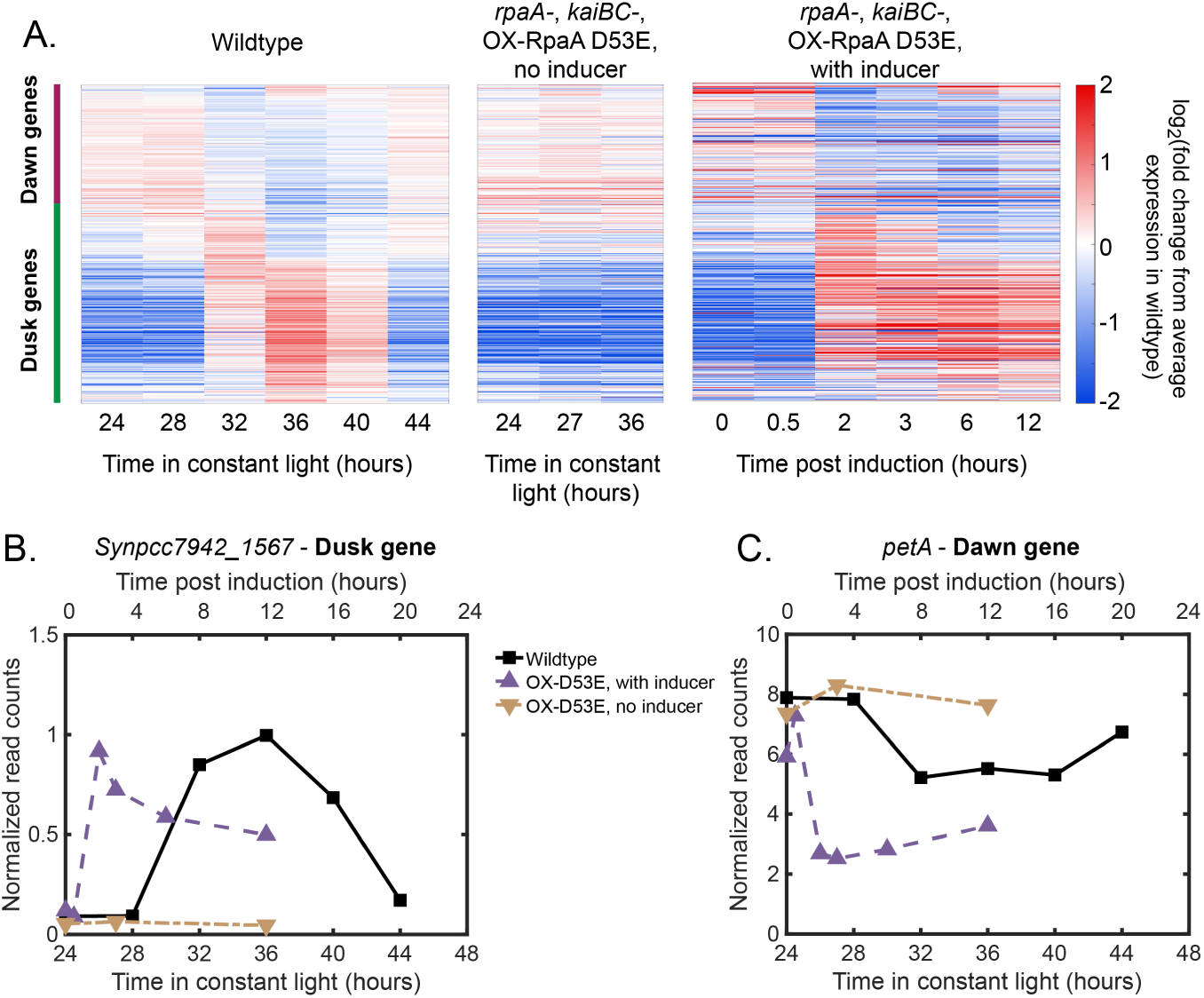
Gene expression dynamics of dusk and dawn circadian genes under Constant Light conditions (data from ref. [10]). **(A)** Gene expression dynamics of circadian genes over 24 hr in Constant Light conditions in wildtype cells (left heat map) and over 12 hr in OX-D53E cells (*rpaA-*, *kaiBC-*, *Ptrc::rpaA(D53E)*) (middle and right heat maps) as measured by RNA sequencing. The OX-D53E strain allows experimental control of RpaA activity via IPTG-inducible expression of the RpaA phosphomimetic RpaA-D53E in cells that lack wildtype RpaA [10]. In the middle panel RpaA-D53E is not induced, and in the right panel IPTG was added to induce RpaA-D53E. Gene expression is quantified as the log_2_ fold change from the average expression of the gene over all time points in the wildtype cells under the Constant Light condition. Genes are ordered based on their phase in Constant Light [8], with the group of dusk and dawn genes indicated with colored bars next to the left heat map. Dusk genes increase in expression under Constant Light in wildtype cells and are maximally expressed when cells expect dusk (left panel, subjective dusk occurs at 36 hours in Constant Light). Note that these dynamics are very similar to the dynamics we observe in Low Light conditions in which the cells periodically experience 12 hr of darkness (Figure 2B, top panel). Dusk genes have constant low expression in OX-D53E cells without inducer which lack RpaA (center) and increase in expression when the RpaA∼P mimic RpaA-D53E is induced in this background (OX-D53E with inducer, right).**(B)** Data from (A) plotted for the representative dusk gene *Synpcc7942*_*1567*. **(C)** Data from (A) plotted for the representative dawn gene *petA*.

**Figure 2-figure supplement 3:**
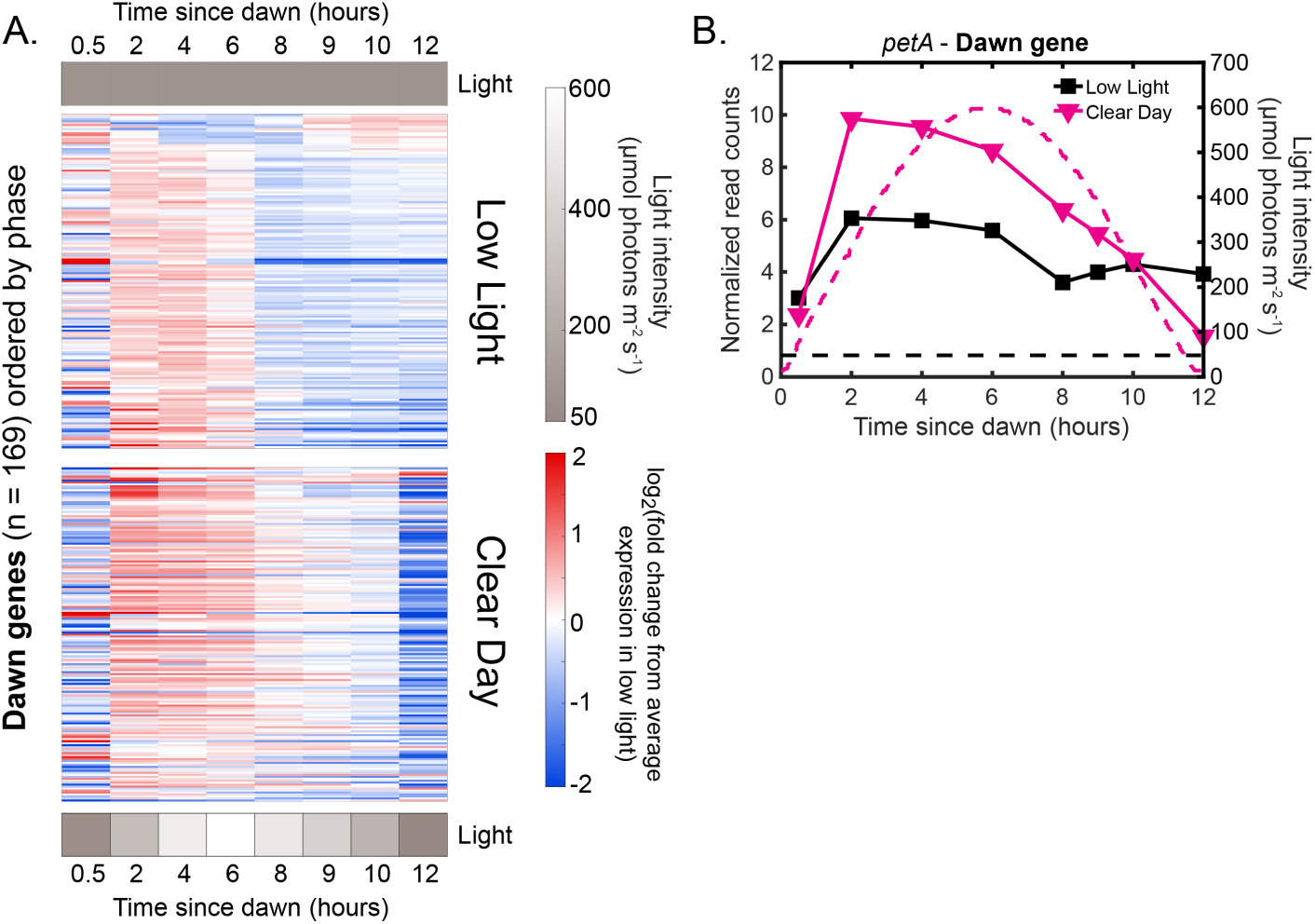
Dawn gene expression increases during the early part of Clear Day relative to Low Light conditions. **(A)** Gene expression dynamics of dawn genes (*n* = 169) under Low Light (top) and Clear Day (bottom) conditions. Gene expression is quantified as the log_2_ fold change from the average expression of the gene over all time points in the Low Light condition (See Materials and Methods - RNA sequencing for more details). Genes are plotted in the same order in both heat maps and were sorted by phase under Constant Light conditions [8]. Most dawn genes are expressed more highly during the early part of Clear Day conditions compared to Low Light conditions. **(B)** Gene expression dynamics of the representative dawn gene *petA* under Low Light (black) and Clear Day (magenta) conditions as measured by RNA sequencing (left y-axis). The light profile for each condition is plotted as dashed lines of the same color with values corresponding to the right y-axis.

**Table S3:** Wiring the AD7376 potentiometer. We used the SOIC-16 housing for the AD7376 potentiometer for ease of soldering to wires. The table indicates how each pin was connected. The length of the GND wire from the Arduino board to the shared ground needs to be kept short (∼2 in. or less) for SPI communication.

| Pin | Connection |
| --- | --- |
| 1 | + of 10 V power supply |
| 2 | GND (shared GND between that of 10V power supply and Arduino) |
| 3 | GND |
| 4 | GND |
| 5 | pin 10 on Arduino (or any other pin designated as a Slave Select, such as 5, 6, or 9) |
| 6 | +5V of Arduino |
| 7 | pin 13 on Arduino (SCLK) |
| 8 | NC (not connected) |
| 9 | NC |
| 10 | NC |
| 11 | pin 11 on Arduino (MOSI) |
| 12 | +5V of Arduino |
| 13 | NC |
| 14 | + of 10V power supply |
| 15 | NC |
| 16 | DIM line of FlexBlock |

**Figure 2-figure supplement 4:**
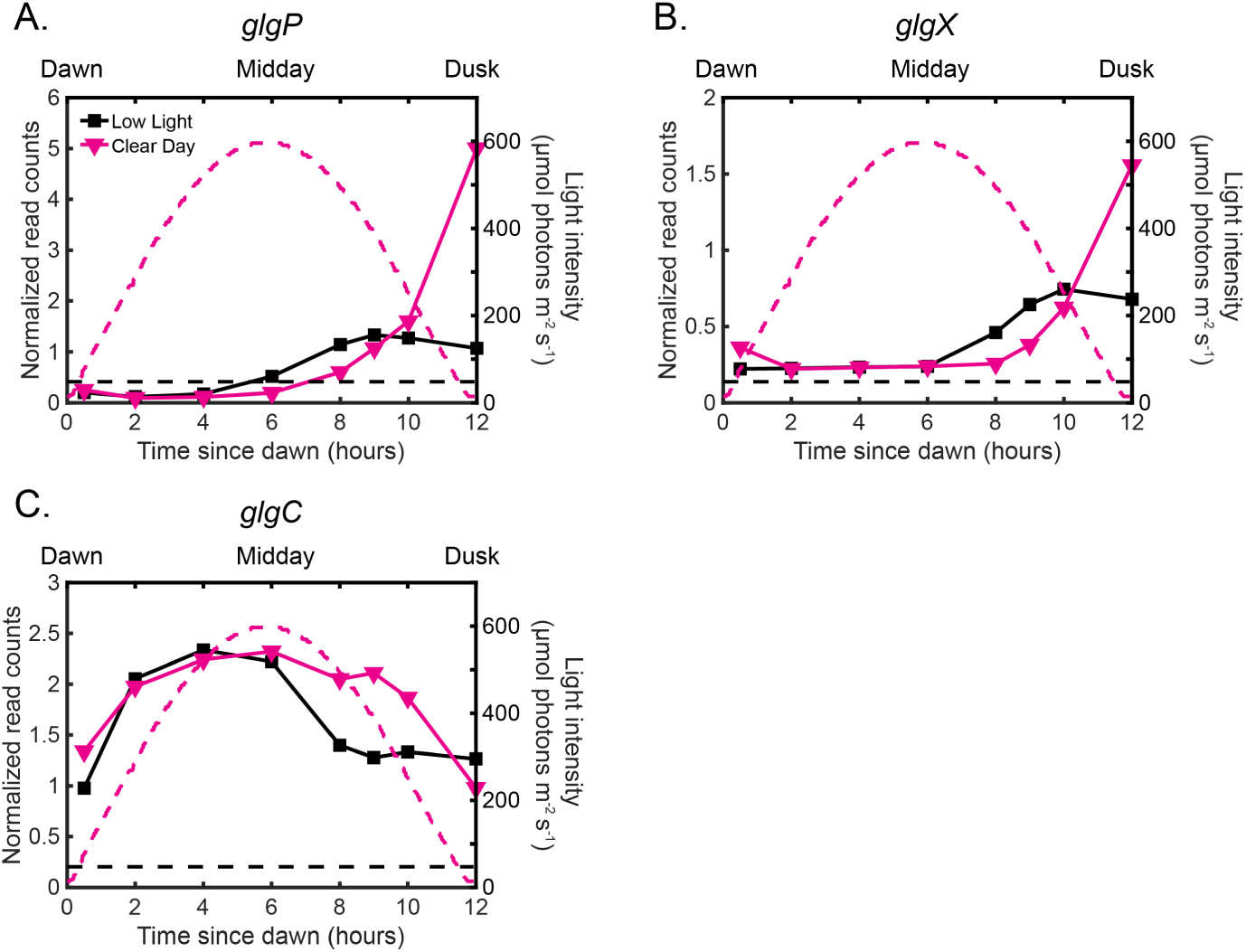
The gene expression dynamics of glycogen production and breakdown enzymes change in Clear Day conditions relative to Low Light conditions. **(A)** Gene expression dynamics of the dusk gene *glgP*, encoding a key enzyme in glycogen breakdown, under Low Light (black) and Clear Day (magenta) conditions as measured by RNA sequencing (left y-axis). The light profile for each condition is plotted as dashed lines of the same color with values corresponding to the right y-axis. **(B)** Gene expression dynamics of the dusk gene *glgX*, a key enzyme in glycogen breakdown, measured and plotted as in (A). **(C)** Gene expression dynamics of the dusk gene *glgC*, a key enzyme in glycogen production, measured and plotted as in (A).

**Table S4:** Fitting bounds. Bounds used for fitting the variables in our simple model of gene expression. H is the Hill coefficient, *β* is the max transcription rate, *α* is the decay/dilution rate, *B* is the background transcription rate, and *K* is a coefficient of activation/repression (see equations 1-3, p. 20-20). The units of *β*, *α*, and *B* are normalized expression/hr; *K* is in normalized expression units.

| Variable | Lower bound | Upper bound |
| --- | --- | --- |
| H | 0 | 7 |
| $\beta$ | 0 | 80 |
| $\alpha$ | 0 | 80 |
| B | 0 | 10 |
| K | 0 | 1 |

**Figure 3-figure supplement 1:**
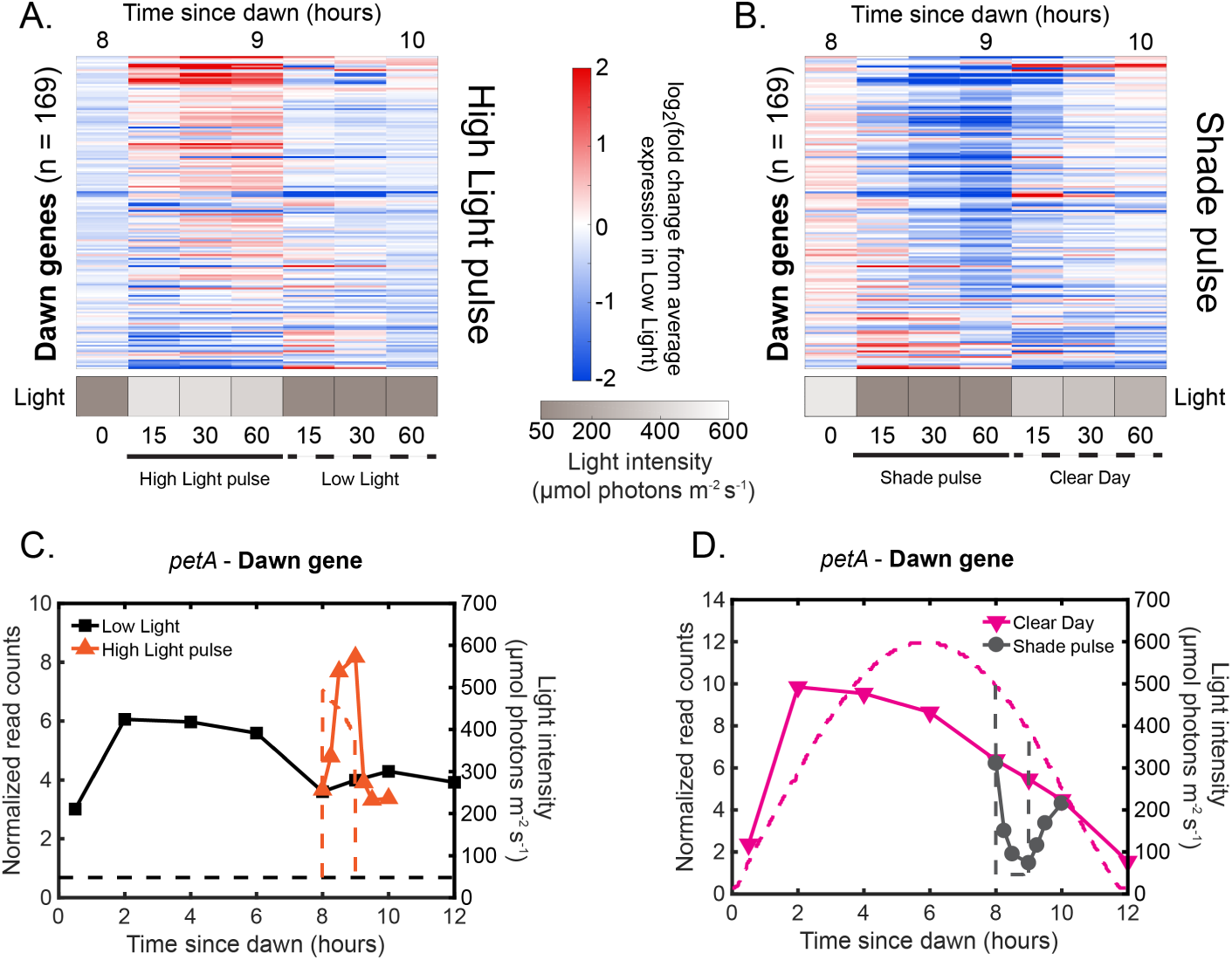
Rapid changes in light intensity affect dawn gene expression in an opposite direction compared to dusk gene expression. **(A)** Gene expression dynamics of dawn genes (*n* = 169) under High Light pulse conditions. Gene expression is quantified as the log_2_ fold change from the average expression of the gene over all time points in the Low Light condition (See Materials and Methods - RNA sequencing for more details). Light intensity at each time point in the High Light pulse condition is indicated in a grayscale heat map. Most dawn genes increase in expression after exposure to the High Light pulse. **(B)** Gene expression dynamics of all dawn genes (*n* = 169) under Shade pulse conditions, plotted as in (A). Genes are ordered the same in **(A)** and (B), sorted by phase under Constant Light conditions [8]. Most dawn genes decrease in expression after exposure to the Shade pulse. **(C)** Gene expression dynamics of the representative dawn gene *petA* under Low Light (black) and High Light pulse (orange) conditions (left y-axis) as measured by RNA sequencing. The light profile for each condition is plotted as dashed lines of the same color with values corresponding to the right y-axis. **(D)** Gene expression dynamics of the representative dawn gene *petA* under Clear Day (magenta) and Shade pulse (gray) conditions, plotted as in (C).

**Figure 3-figure supplement 2:**
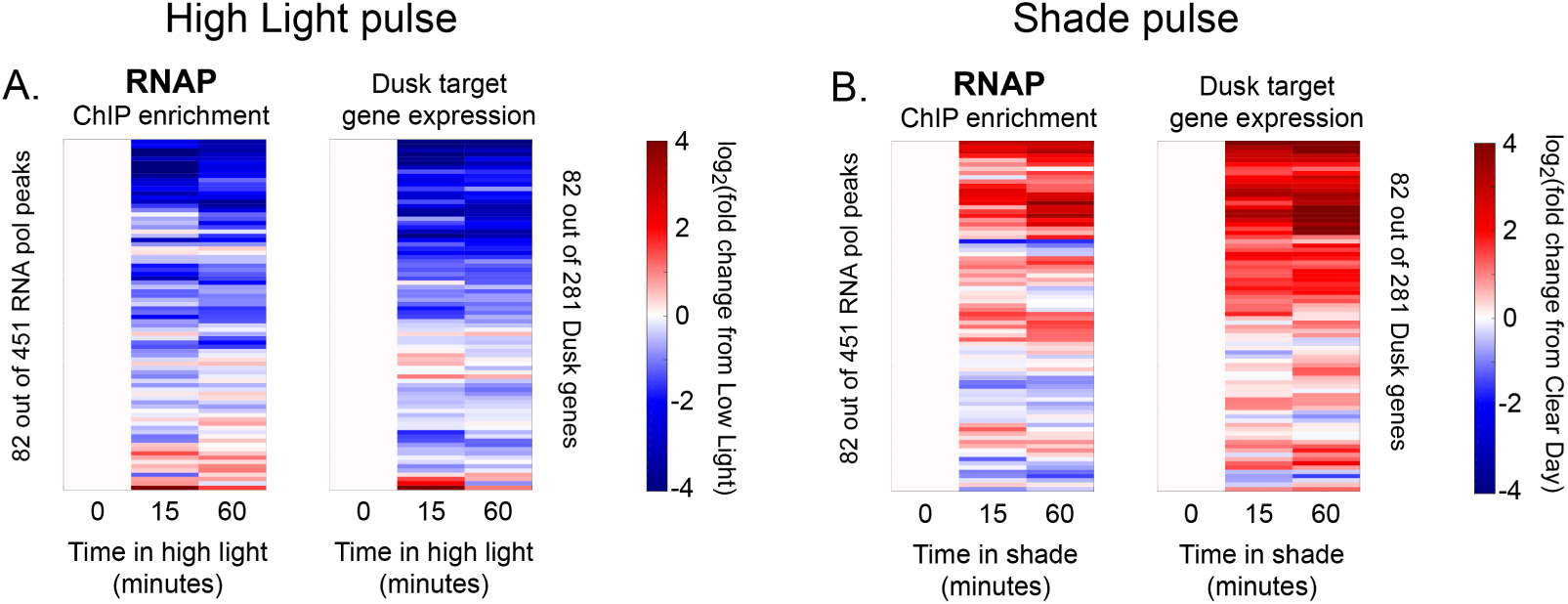
Changes in RNAP enrichment and downstream dusk gene expression after rapid changes in light intensity. **(A)** Changes in enrichment of RNAP upstream of dusk genes during High Light pulse conditions (left heat map) and corresponding changes in target dusk gene expression (right heat map) for the 82 dusk genes with RNAP peaks in their promoters. ChIP enrichment (left heat map) is quantified as the log_2_ fold change of enrichment in High Light compared to enrichment at 8 hours since dawn in Low Light conditions (time zero). The right heat map shows the change in expression of the gene target of the corresponding RNAP peak expressed as the log_2_ fold change of expression in High Light compared to expression at 8 hours since dawn in Low Light conditions (time zero). An RNAP peak and its target gene are aligned horizontally in the two heat maps so that they can be directly compared side-by-side. See Materials and Methods, ChIP-seq analysis for more details. **(B)** Changes in enrichment of RNAP upstream of dusk genes during Shade pulse conditions (left heat map) and corresponding changes in target dusk gene expression (right heat map). ChIP enrichment (left heat map) is quantified as the log_2_ fold change of enrichment in Shade from enrichment at 8 hours since dawn in Clear Day conditions (time zero). The right heat map shows the change in expression of the gene target of the corresponding RNAP peak expressed as the log_2_ fold change of expression in Shade from expression at 8 hours since dawn in Clear Day conditions (time zero). An RNAP peak and its target gene are aligned in the two heat maps. RNAP peaks and genes have the same order in **(A)** and(B). See Materials and Methods, ChIP-seq analysis for more details. The correlation between RNAP enrichment change and downstream dusk gene expression reported in Figure 3G also holds after 15 minutes of exposure to High Light or Shade (High Light correlation = .6386, Shade correlation = .6806, changes after 15 minutes).

**Figure 4-figure supplement 1:**
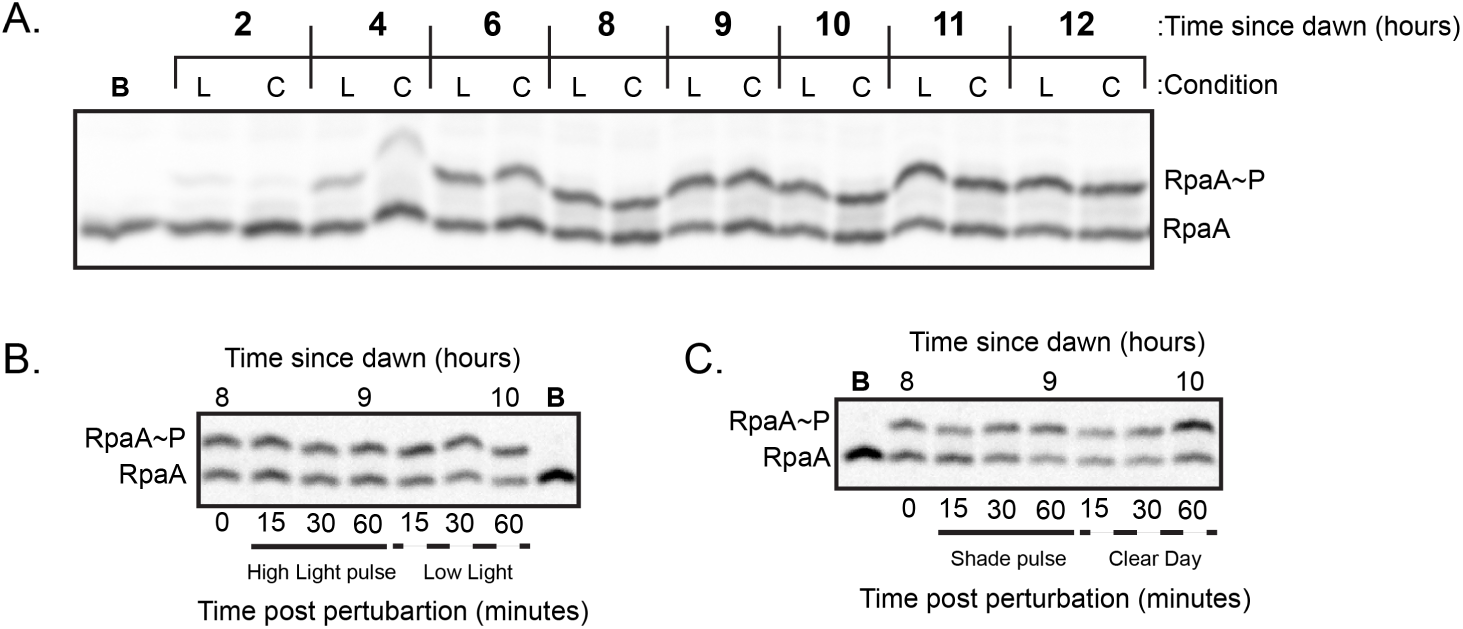
Representative Western blots used to quantify relative levels of RpaA∼P under dynamic light conditions. **(A)** Representative Western Blot used to quantify levels of RpaA∼P under Low Light and Clear Day conditions. Lysates were prepared from cells harvested from either Low Light (L) or Clear Day **(C)** conditions at the indicated time and subject to Phos-tag electrophoresis and Western blotting with an anti-RpaA antibody (see Methods). One sample was boiled prior to loading (Lane indicated with ‘B’) to identify the heat-labile band on the gel corresponding to RpaA∼P. **(B)** Representative Western Blot used to quantify levels of RpaA∼P under High Light pulse conditions. Time 0 refers to 8 hours since dawn under Low Light conditions. **(C)** Representative Western Blot used to quantify levels of RpaA∼P under Shade pulse conditions. Time 0 refers to 8 hours since dawn under Clear Day conditions.

**Figure 4-figure supplement 2:**
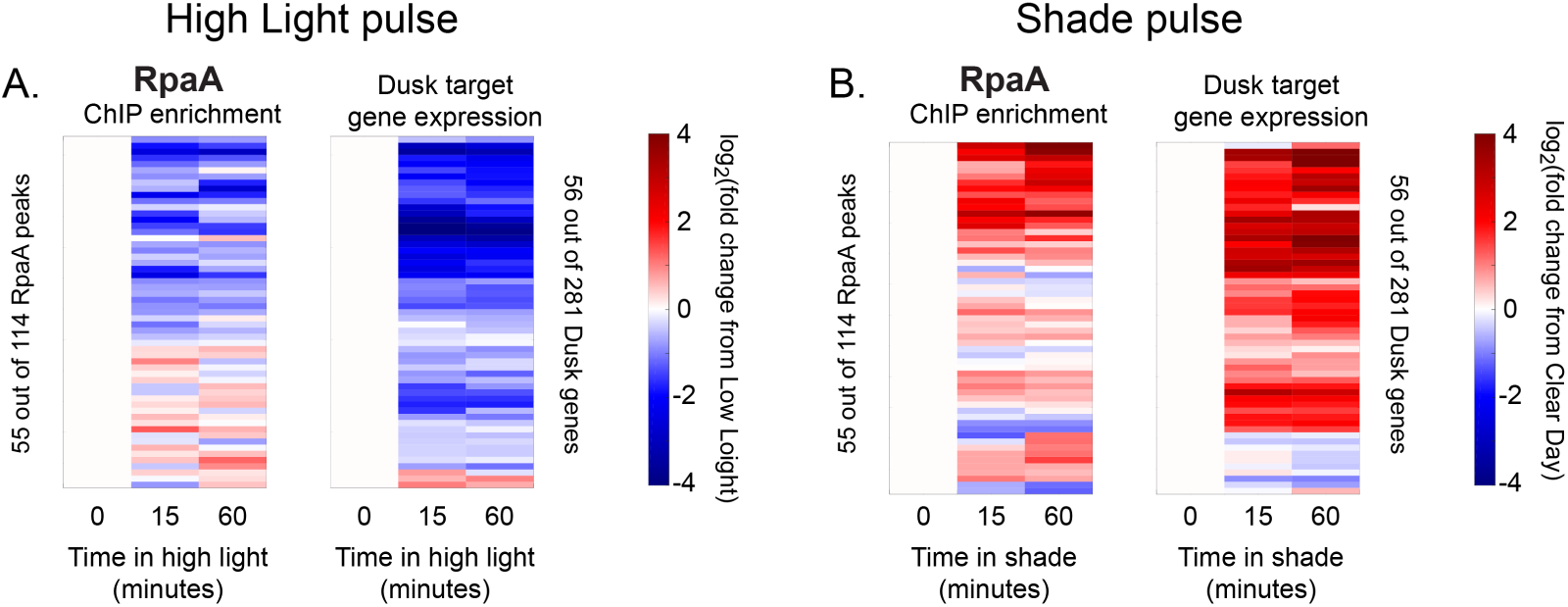
Changes in RpaA enrichment and downstream dusk gene expression after rapid changes in light intensity. **(A)** Changes in enrichment of RpaA upstream of dusk genes during High Light pulse conditions (left heat map) and corresponding changes in target dusk gene expression (right heat map) for the 56 dusk genes with RpaA peaks in their promoters. ChIP enrichment (left heat map) is expressed as the log_2_ fold change of enrichment in High Light relative to enrichment at 8 hours since dawn in Low Light conditions (time zero). The right heat map shows the change in expression of the gene target of the corresponding RpaA peak expressed as the log_2_ fold change of expression in High Light from expression at 8 hours since dawn in Low Light conditions (time zero). An RpaA peak and its target gene are aligned horizontally in the two heat maps. See Materials and Methods, ChIP-seq analysis for more details. **(B)** Changes in enrichment of RpaA upstream of dusk genes during Shade pulse conditions (left heat map) and corresponding changes in target dusk gene expression (right heat map). ChIP enrichment (left heat map) is expressed as the log_2_ fold change of enrichment in Shade relative to enrichment at 8 hours since dawn in Clear Day conditions (time zero). The right heat map shows the change in expression of the gene target of the corresponding RpaA peak expressed as the log_2_ fold change of expression in Shade from expression at 8 hours since dawn in Clear Day conditions (time zero). An RpaA peak and its target gene are aligned horizontally in the two heat maps. Peaks and genes have the same order in **(A)** and (B). See Materials and Methods, ChIP-seq analysis for more details. The correlation between RpaA change and downstream dusk gene expression reported in Figure 4C also holds after 15 minutes of exposure to High Light or Shade (High Light correlation = .5433, Shade correlation = .3808, changes after 15 minutes).

**Figure 4-figure supplement 3:**
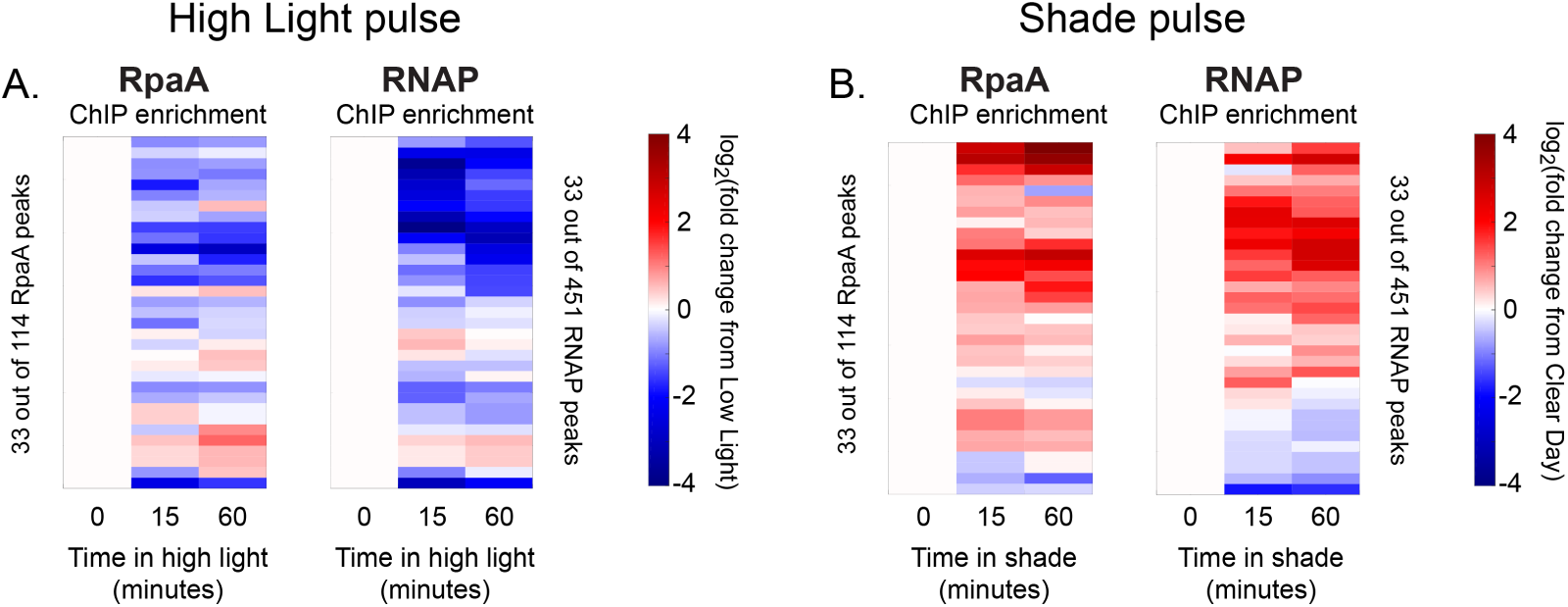
Changes in RpaA and RNA polymerase enrichment upstream of dusk genes after rapid changes in light intensity. **(A)** Changes in enrichment of RpaA upstream of dusk genes during High Light pulse conditions (left heat map) and corresponding changes in RNAP enrichment upstream of the same gene (right heat map) for the 33 dusk genes with RpaA and RNAP peaks in their promoters. ChIP enrichment is quantified as the log_2_ fold change of enrichment in High Light from enrichment at 8 hours since dawn in Low Light conditions (time zero). RpaA and RNAP peaks upstream of the same dusk gene are aligned horizontally in the two heat maps. See Materials and Methods, ChIP-seq analysis for more details. **(B)** Changes in enrichment of RpaA upstream of dusk genes during Shade pulse conditions (left heat map) and corresponding changes in RNAP enrichment upstream of the same gene (right heat map). ChIP enrichment is quantified as the log_2_ fold change of enrichment in Shade from enrichment at 8 hours since dawn in Clear Day conditions (time zero). RpaA and RNAP peaks upstream of the same dusk gene are aligned horizontally in the two heat maps. Peaks have the same order in (A) and (B). See Materials and Methods, ChIP-seq analysis for more details. The correlation between RpaA and RNAP enrichment change upstream of dusk gene expressions reported in Figure 4D also holds after 15 minutes of exposure to High Light or Shade (High Light correlation = .5598, Shade correlation = .4179).

**Figure 4-figure supplement 4:**
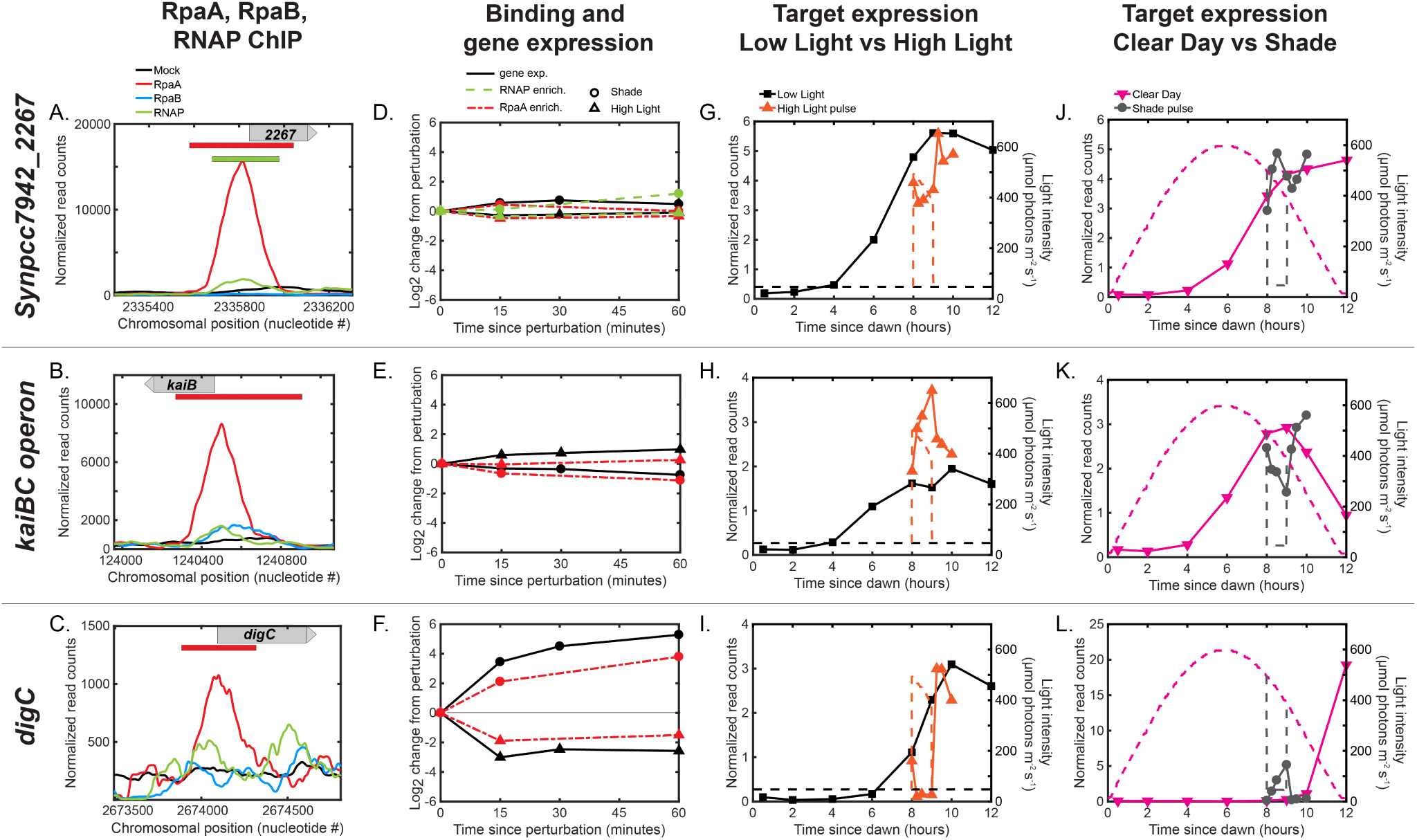
Multifactorial behavior of RpaA∼P at select promoters under changes in light intensity. (A)-**(C)** Normalized ChIP-seq signal of RpaA (red), RpaB (blue), RNAP (green) and mock IP (black) upstream of the **(A)** the representative dusk gene *Synpcc7942 2267*, **(B)** the *kaiBC* operon and **(C)** another representative dusk gene, *digC*, at 8 hours since dawn in Low Light. The chromosomal position of the gene is located on the plot with a gray bar with an arrow indicating directionality of the gene. The location of RpaA and RNAP peaks are indicated on top of the plot with red (RpaA) and green (RNAP) bars. No RpaB peaks were found upstream of these genes. No RNAP peak was found upstream of *kaiB* or *digC*. See Materials and Methods, ChIP-seq analysis for more details. **(D)**-**(F)** Changes in enrichment of RpaA (red) and RNAP (green) and downstream gene expression (black) after exposure to the High Light pulse (triangles) or the Shade pulse (circles) for (D) *Synpcc7942_2267*, (E) the *kaiBC* operon, and (F) *digC*. See Materials and Methods, ChIP-seq analysis for more details. **(G)**-**(L)** Gene expression dynamics of *Synpcc7942_2267* (G,J) *kaiB* (H,K) and *digC* (I,L) under Low Light vs High Light pulse (G-I) and Clear Day vs Shade pulse (J-L) conditions. RpaA binding does not change upstream of the dusk genes *Synpcc7942_2267* and *kaiB* after changes in light (D,E), and the expression of these genes does not change significantly in response to changes in light intensity (G,J;H,K). In contrast, RpaA binding changes significantly (F) upstream of the light-responsive dusk gene *digC* (I,L).

**Figure 5-figure supplement 1:**
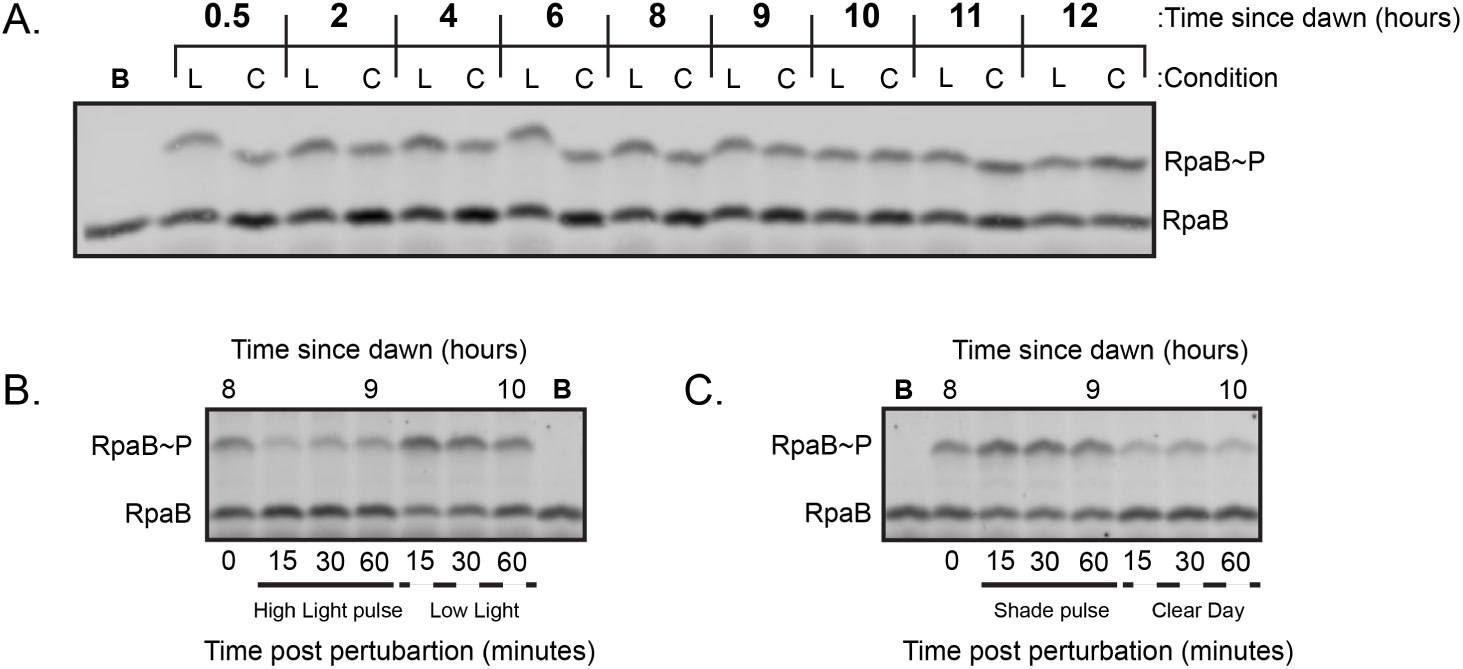
Representative Western blots used to quantify relative levels of RpaB∼P under dynamic light conditions. **(A)** Representative Western Blot used to quantify levels of RpaB∼P under Low Light and Clear Day conditions. Lysates were prepared from cells harvested from either Low Light (L) or Clear Day **(C)** conditions at the indicated time and subject to Phos-tag electrophoresis and Western blotting with an anti-RpaB antibody (see Methods). One sample was boiled prior to loading (Lane indicated with ‘B’) to identify the heat-labile band on the gel corresponding to RpaB∼P. **(B)** Representative Western Blot used to quantify levels of RpaB∼P under High Light pulse conditions. Time 0 refers to 8 hours since dawn under Low Light conditions. **(C)** Representative Western Blot used to quantify levels of RpaB∼P under Shade pulse conditions. Time 0 refers to 8 hours since dawn under Clear Day conditions.

**Figure 5-figure supplement 2:**
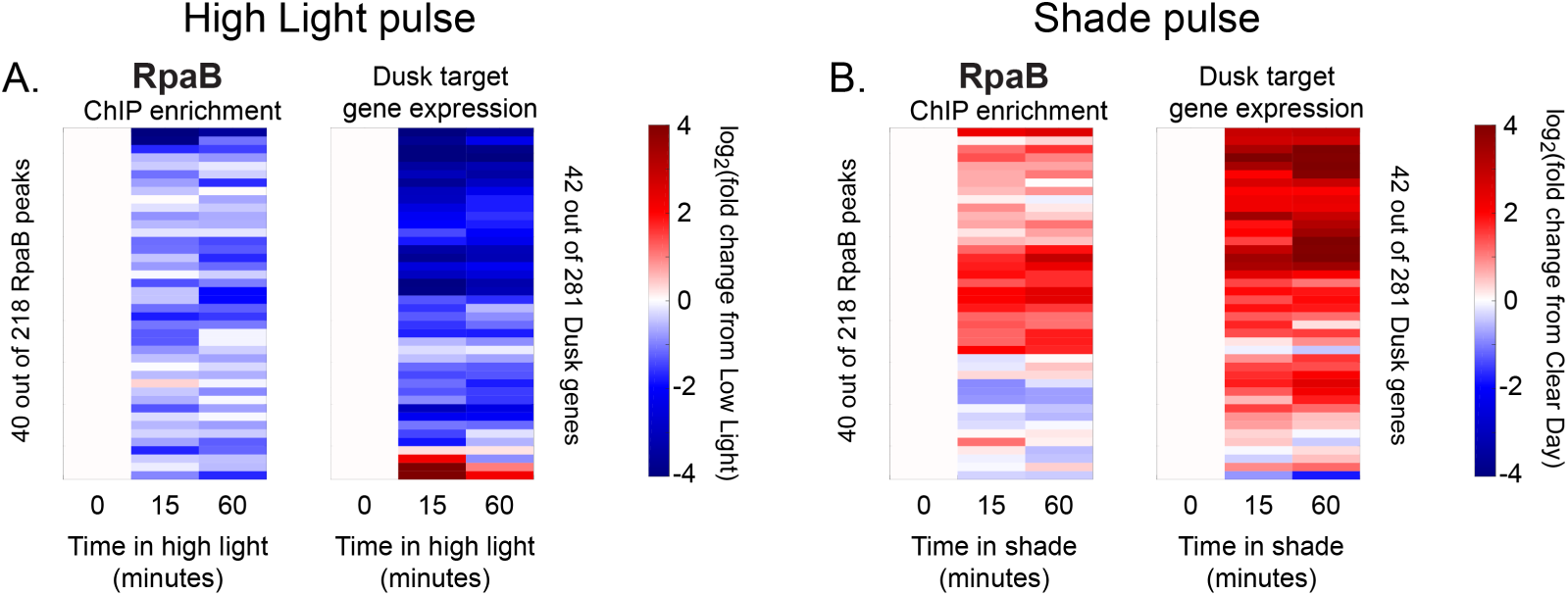
Changes in RpaB enrichment and downstream dusk gene expression after rapid changes in light intensity. **(A)** Changes in enrichment of RpaB upstream of dusk genes during High Light pulse conditions (left heat map) and corresponding changes in target dusk gene expression (right heat map) for the 42 dusk genes with RpaB peaks in their promoters. ChIP enrichment (left heat map) is quantified as the log_2_ fold change of enrichment in High Light from enrichment at 8 hours since dawn in Low Light conditions (time zero). The right heat map shows the change in expression of the gene target of the corresponding RpaB peak quantified as the log_2_ fold change of expression in High Light from expression at 8 hours since dawn in Low Light conditions (time zero). An RpaB peak and its target gene are aligned horizontally in the two heat maps. See Materials and Methods, ChIP-seq analysis for more details. **(B)** Changes in enrichment of RpaB upstream of dusk genes during Shade pulse conditions (left heat map) and corresponding changes in target dusk gene expression (right heat map). ChIP enrichment (left heat map) is quantified as the log_2_ fold change of enrichment in Shade from enrichment at 8 hours since dawn in Clear Day conditions (time zero). The right heat map shows the change in expression of the gene target of the corresponding RpaB peak quantified as the log_2_ fold change of expression in Shade from expression at 8 hours since dawn in Clear Day conditions (time zero). An RpaB peak and its target gene are aligned horizontally in the two heat maps. Peaks and genes have the same order in (A) and (B). See Materials and Methods, ChIP-seq analysis for more details. The correlation between RpaB change and downstream dusk gene expression reported in Figure 5C also holds after 15 minutes of exposure to High Light or Shade (High Light correlation = .1637, Shade correlation = .3520, changes after 15 minutes).

**Figure 5-figure supplement 3:**
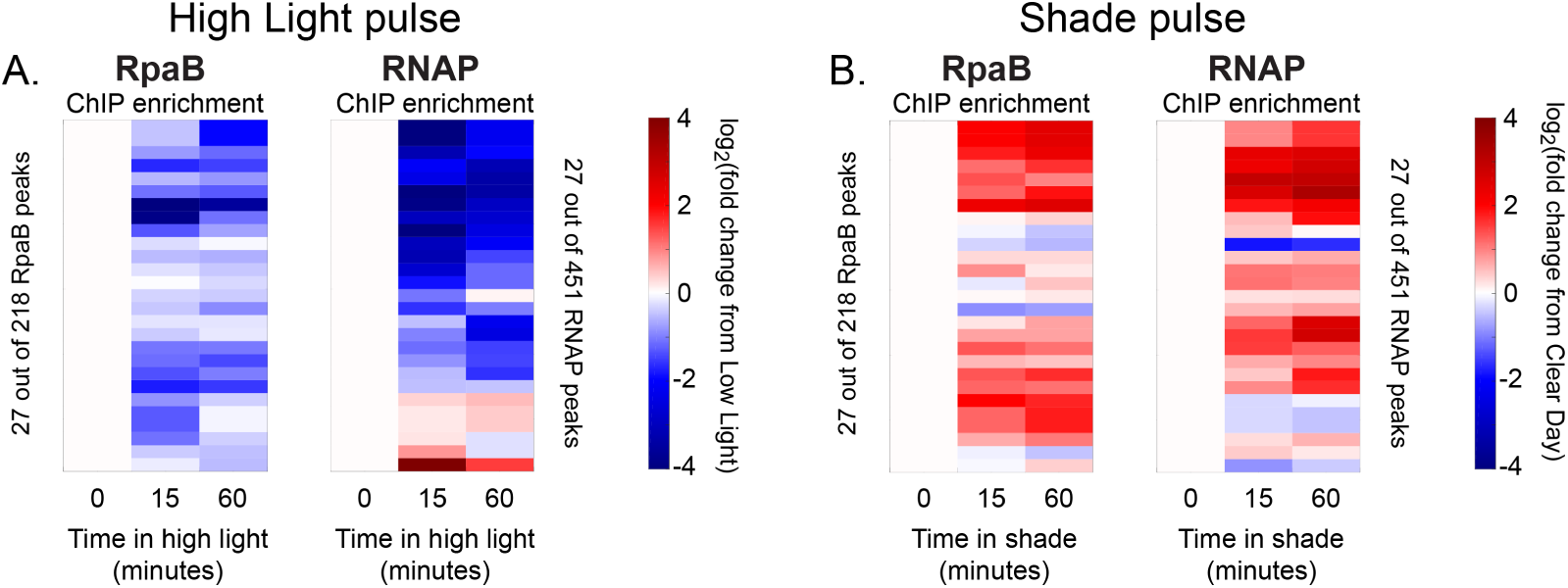
Changes in RpaB and RNA polymerase enrichment upstream of dusk genes after rapid changes in light intensity. **(A)** Changes in enrichment of RpaB upstream of dusk genes during High Light pulse conditions (left heat map) and corresponding changes in RNAP enrichment upstream of the same gene (right heat map) for the 27 dusk genes with RpaB and RNAP peaks in their promoters. ChIP enrichment is expressed as the log_2_ fold change of enrichment in High Light from enrichment at 8 hours since dawn in Low Light conditions (time zero). RpaB and RNAP peaks upstream of the same dusk gene are aligned horizontally in the two heat maps. See Materials and Methods, ChIP-seq analysis for more details. **(B)** Changes in enrichment of RpaB upstream of dusk genes during Shade pulse conditions (left heat map) and corresponding changes in RNAP enrichment upstream of the same gene (right heat map). ChIP enrichment is expressed as the log_2_ fold change of enrichment in Shade from enrichment at 8 hours since dawn in Clear Day conditions (time zero). RpaB RNAP peaks upstream of the same dusk gene are aligned horizontally in the two heat maps. Peaks have the same order in (A) and (B). See Materials and Methods, ChIP-seq analysis for more details. The correlation between RpaA and RNAP enrichment change upstream of dusk gene expressions reported in Figure 5D also holds after 15 minutes of exposure to High Light or Shade (High Light correlation = .2185, Shade correlation = .4151).

**Table S5:**
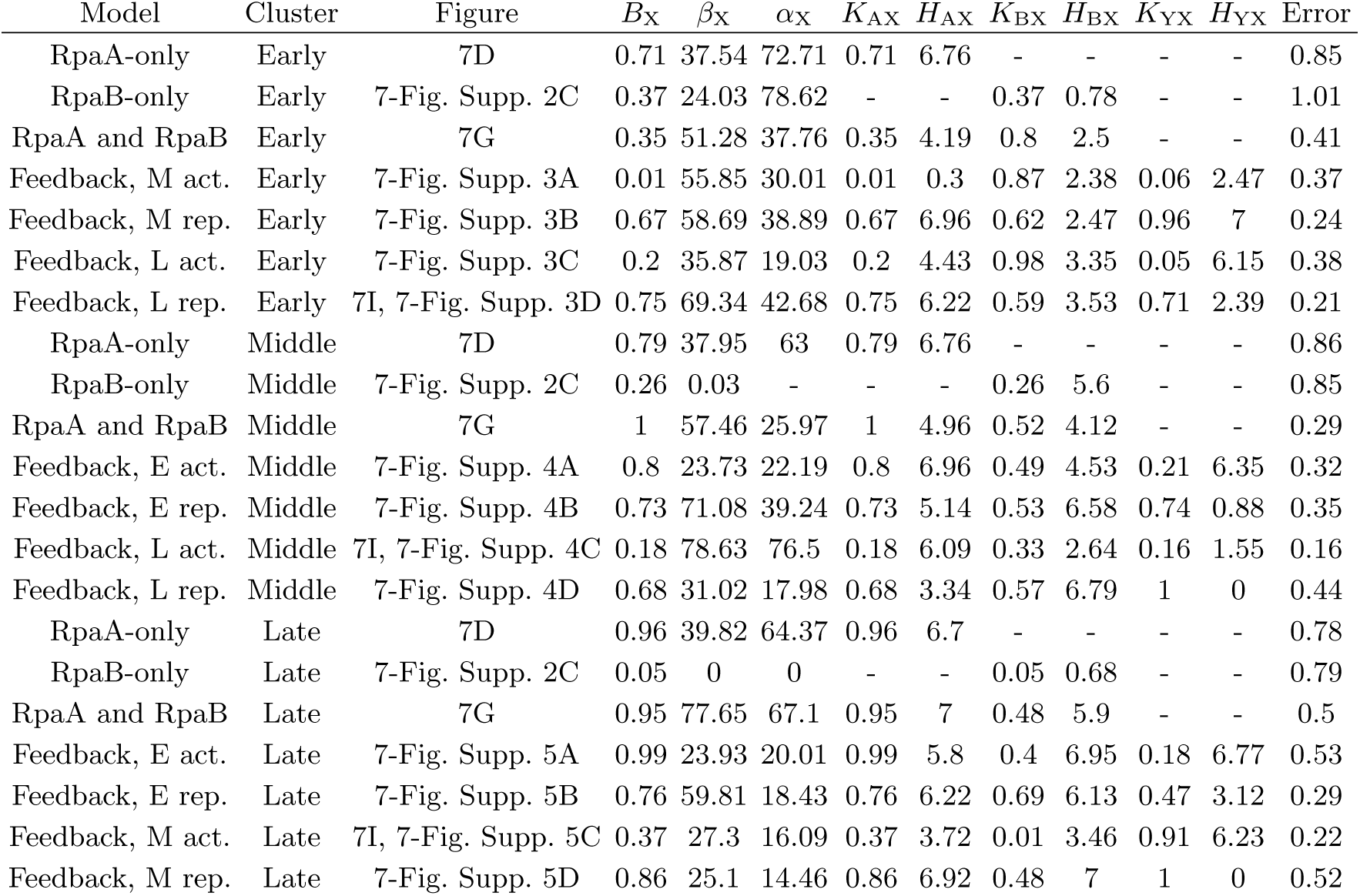
Fitting results. The definitions of the variables are given in equations 1-3, p. 20-20. The error is defined as the square root of the sum of the squared deviations between simulation and data.

**Figure 6-figure supplement 1:**
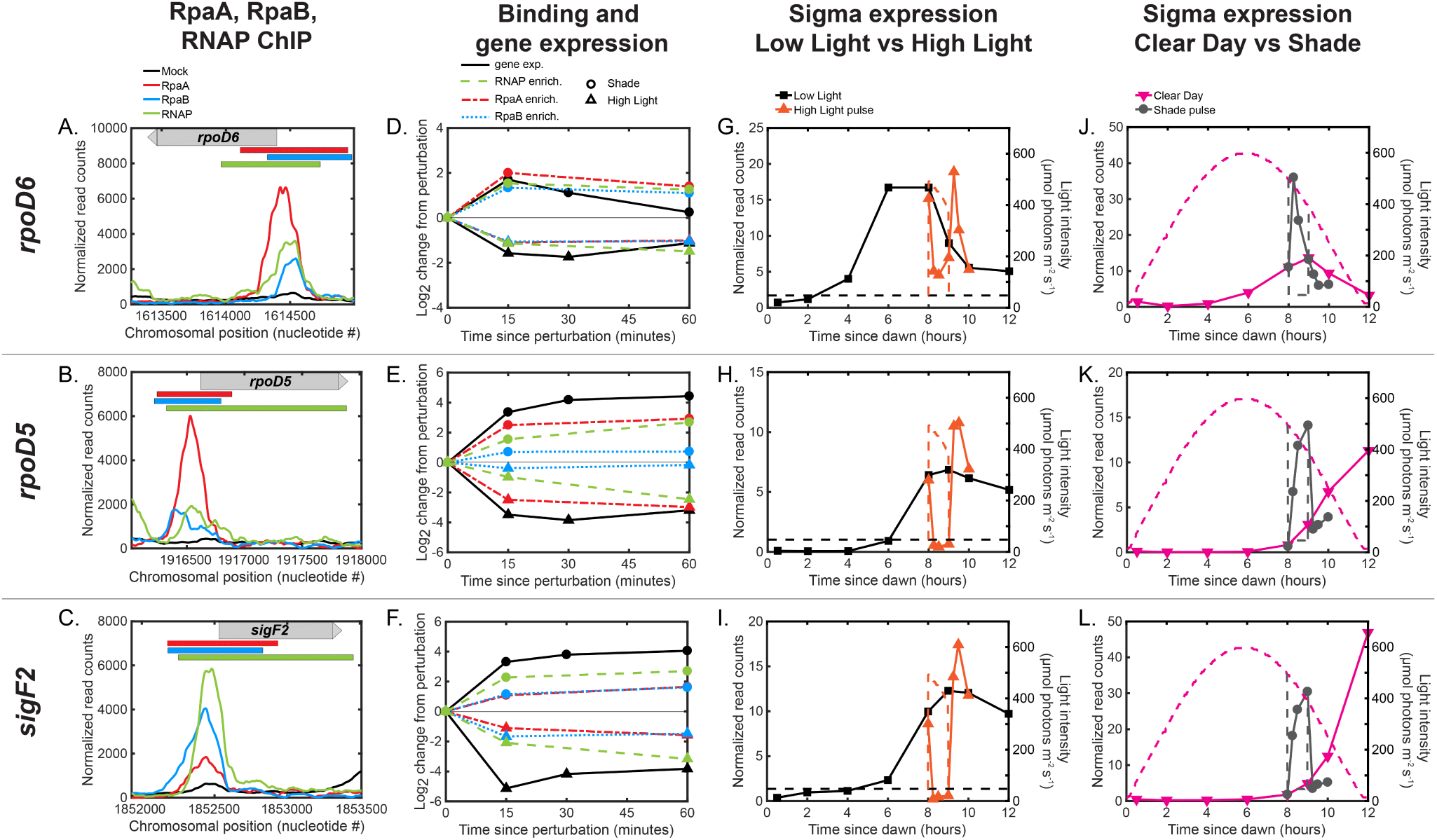
Regulation of dusk sigma factor gene expression by RpaA and RpaB. **(A)-(C)** Normalized ChIP-seq signal of RpaA (red), RpaB (blue), RNAP (green) and mock IP (black) upstream of the sigma factor genes (A) *rpoD6*, (B) *rpoD5*, and **(C)** *sigF2*. The location of the gene is located on the plot with a gray bar with an arrow indicating directionality of the gene. The location of RpaA, RpaB, and RNAP peaks are indicated on top of the plot with red (RpaA), blue (RpaB), and green (RNAP) bars. See Materials and Methods, ChIP-seq analysis for more details. (D)-**(F)** Changes in enrichment of RpaA (red), RpaB (blue), and RNAP (green) and downstream sigma factor gene expression (black) after exposure to the High Light pulse (triangles) or the Shade pulse (circles) upstream of *rpoD6* (D), *rpoD5* (E), and *sigF2*(F). See Materials and Methods, ChIP-seq analysis for more details. **(G)**-**(L)** Gene expression dynamics of *rpoD6* (G,J), *rpoD5* (H,K), and *sigF2* (I,L) under Low Light vs High Light pulse **(G)-(I)** and Clear Day vs Shade pulse **(J)-(L)** conditions. RpaA and RpaB binding changes in a correlated manner upstream of these genes. RpaA and RpaB binding also correlates with changes in RNAP enrichment and sigma factor expression levels.

**Figure 7-figure supplement 1:**
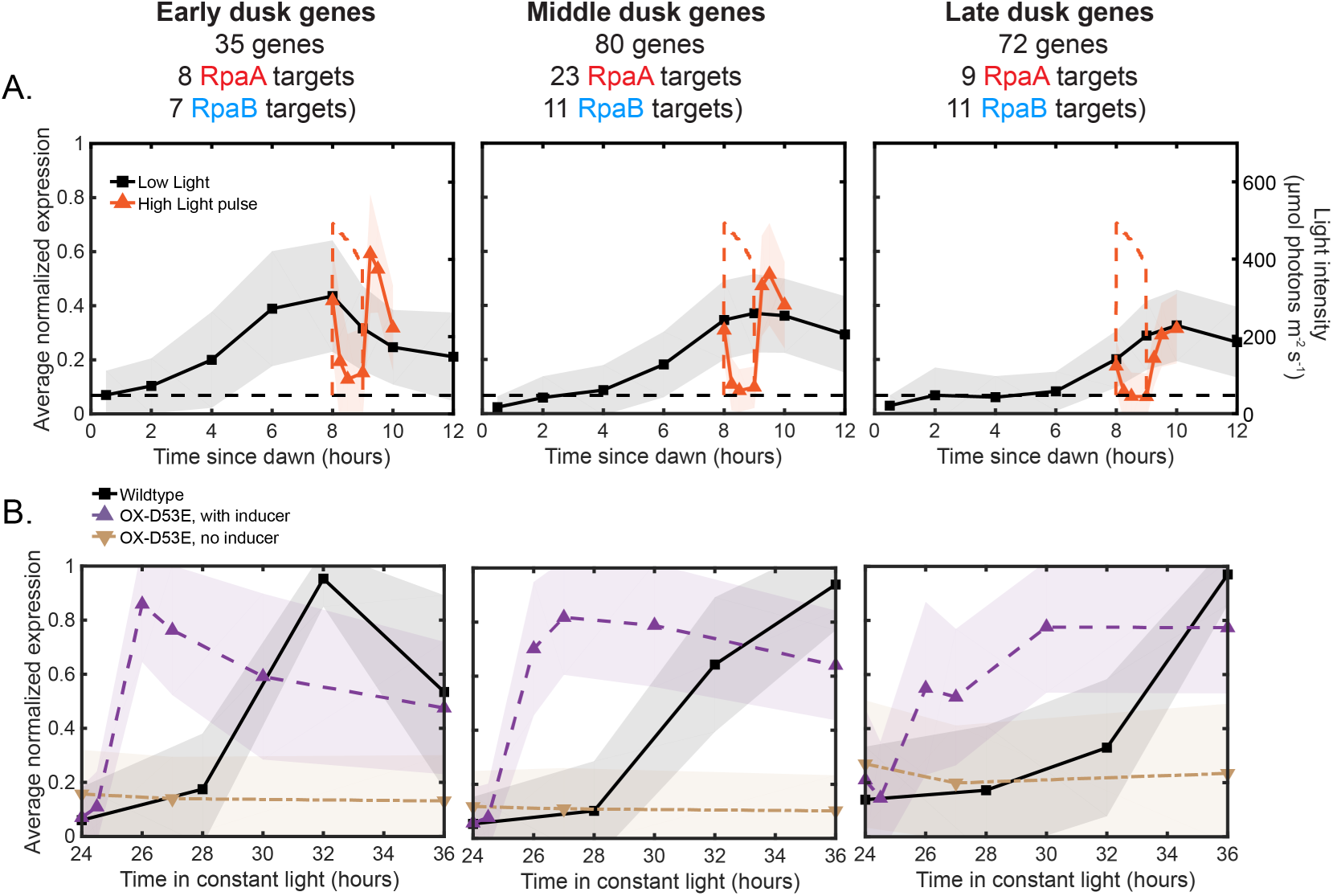
Average expression profiles of the major dusk gene clusters under various conditions. **(A)** Average expression profiles of the Early (left plot), Middle (middle plot), and Late (right plot) dusk gene clusters under Low Light (black) and High Light pulse (orange) conditions (left y-axis). The expression values of each gene across all 4 light conditions in this work were normalized to a range of 0 to 1, and the normalized expression values were averaged within each cluster. The shaded region of the plot indicates the standard deviation of the normalized expression values within the cluster. Lists of genes belonging to each cluster and the scaled expression values are available in Figure 7-source data 1. The light intensity profile for each condition is plotted as dashed lines in the same color with values corresponding to the right y-axis. **(B)** Average expression profiles of the Early (left plot), Middle (middle plot), and Late (right plot) dusk gene clusters in Constant Light conditions in wildtype and OX-D53E cells (*rpaA-*, *kaiBC-*, *Ptrc::rpaA(D53E)*) (data from [10]). The OX-D53E strain allows experimental control of RpaA activity via IPTG-inducible expression of the RpaA phosphomimetic RpaA-D53E in cells that lack wildtype RpaA. Plotted are average cluster expression in wildtype cells in Constant Light conditions (black squares), OX-D53E cells without inducer (RpaA phosphomimetic not induced, brown downward triangles), and OX-D53E cells with inducer (RpaA phosphomimetic induced, purple upward triangles). The expression values of each gene within each strain in Constant Light (wildtype or OX-D53E) were separately normalized to a range of 0 to 1, and the normalized expression values were averaged within each cluster. Lists of genes belonging to each cluster and the scaled expression values are available in Figure 7-source data 1. The shaded region on the plot indicates the standard deviation of the normalized expression values within the cluster.

**Figure 7-figure supplement 2:**
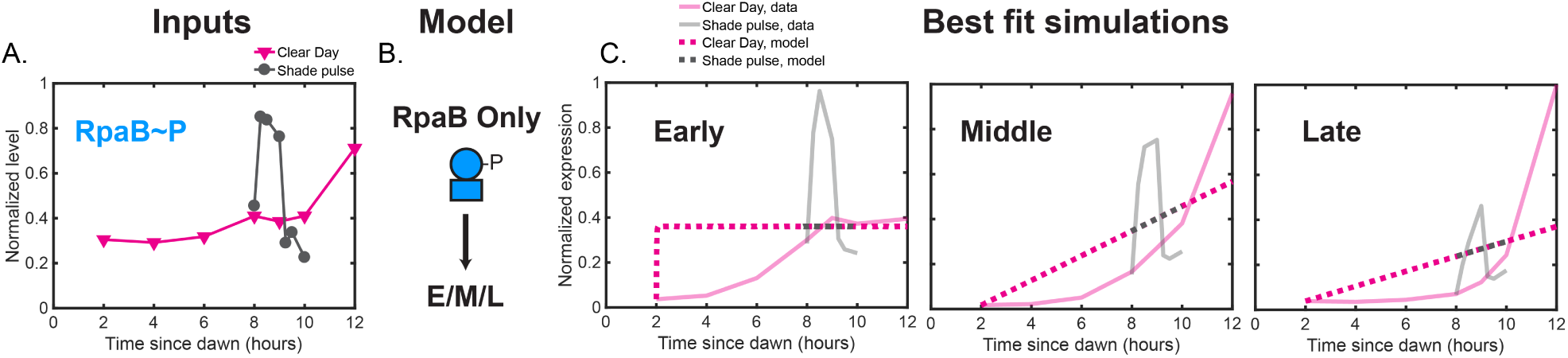
Best fit simulations of ‘RpaB-only’ models in which RpaB∼P solely activates the expression of the dusk gene clusters. **(A)** Normalized RpaB∼P levels under Clear Day (magenta) and Shade pulse (gray) conditions used as model input. **(B)** Model schematic. Dusk gene expression under Clear Day and Shade pulse conditions was modelled as an activation Hill function of RpaB∼P levels only. **(C)** Best fit simulations of RpaB-only models for Early (left plot), Middle (middle plot), and Late (right plot) dusk gene clusters. The average expression values of the cluster used for model fitting are plotted as solid transparent lines, and the simulated data of the best fit model is plotted as dotted lines. Data for Clear Day conditions are plotted in magenta, and Shade pulse in gray.

**Figure 7-figure supplement 3:**
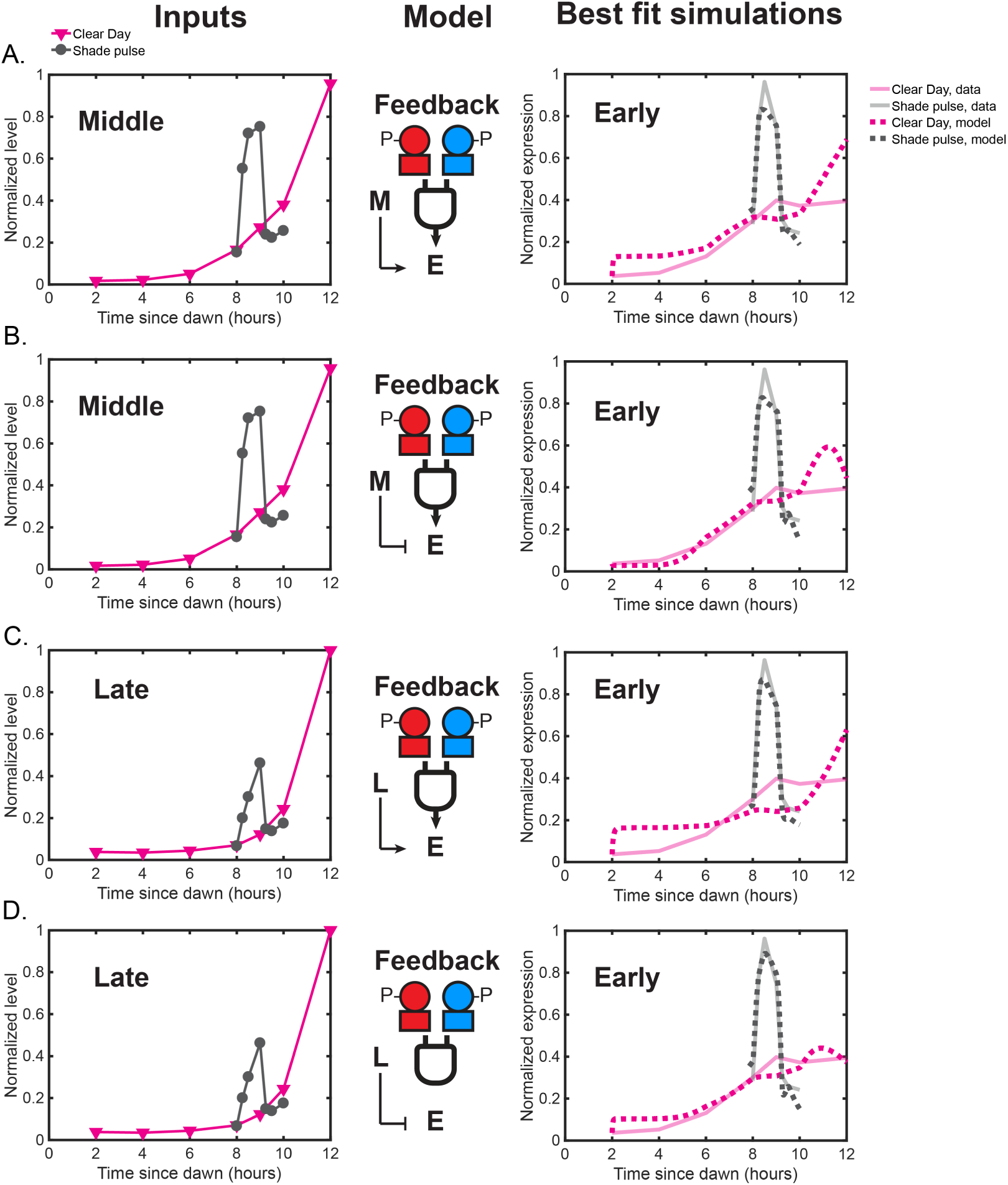
Models in which either the Middle or Late cluster feeds back to influence Early cluster expression. **(A)** Feedback model in which the expression of the Early dusk cluster is an activation Hill function of Middle gene expression and an activation Hill function of both RpaA∼P and RpaB∼P. The left plot shows normalized Middle cluster expression levels under Clear Day (magenta) and Shade pulse (gray) conditions used as model input in addition to the RpaA∼P and RpaB∼P dynamics shown in Figure 7B,E of the main text. The right plot shows the average expression values of the Early cluster data (solid transparent lines), and the simulation produced the best fit model (dotted lines). Data for Clear Day conditions are plotted in magenta, and Shade pulse in gray. **(B)** Feedback model in which Early gene cluster expression is a repression Hill function of Middle cluster expression levels and an activation Hill function of both RpaA∼P and RpaB∼P, presented as in (A). **(C)** Feedback model in which Early cluster expression is an activation Hill function of Late cluster expression levels and an activation Hill function of both RpaA∼P and RpaB∼P. The left plot shows normalized Late cluster expression levels under Clear Day (magenta) and Shade pulse (gray) conditions used as model input. The right plot shows the average expression values of the Early cluster data (solid transparent lines), and the simulation produced the best fit model (dotted lines). Data for Clear Day conditions are plotted in magenta, and Shade pulse in gray. **(D)** Feedback model in which Early cluster expression is a repression Hill function of Late cluster expression levels and an activation Hill function of both RpaA∼P and RpaB∼P, presented as in (C). A model with an incoherent feedforward architecture in which the Late cluster represses Early cluster expression (D) best recapitulates the difference of Early cluster responses to Shade and Clear Day Sunset. During the Shade pulse, the Late cluster levels do not reach high enough levels to inhibit Early cluster expression, but at Sunset in Clear Day, Late cluster levels reach high enough levels to repress the expression of the Early cluster.

**Figure 7-figure supplement 4:**
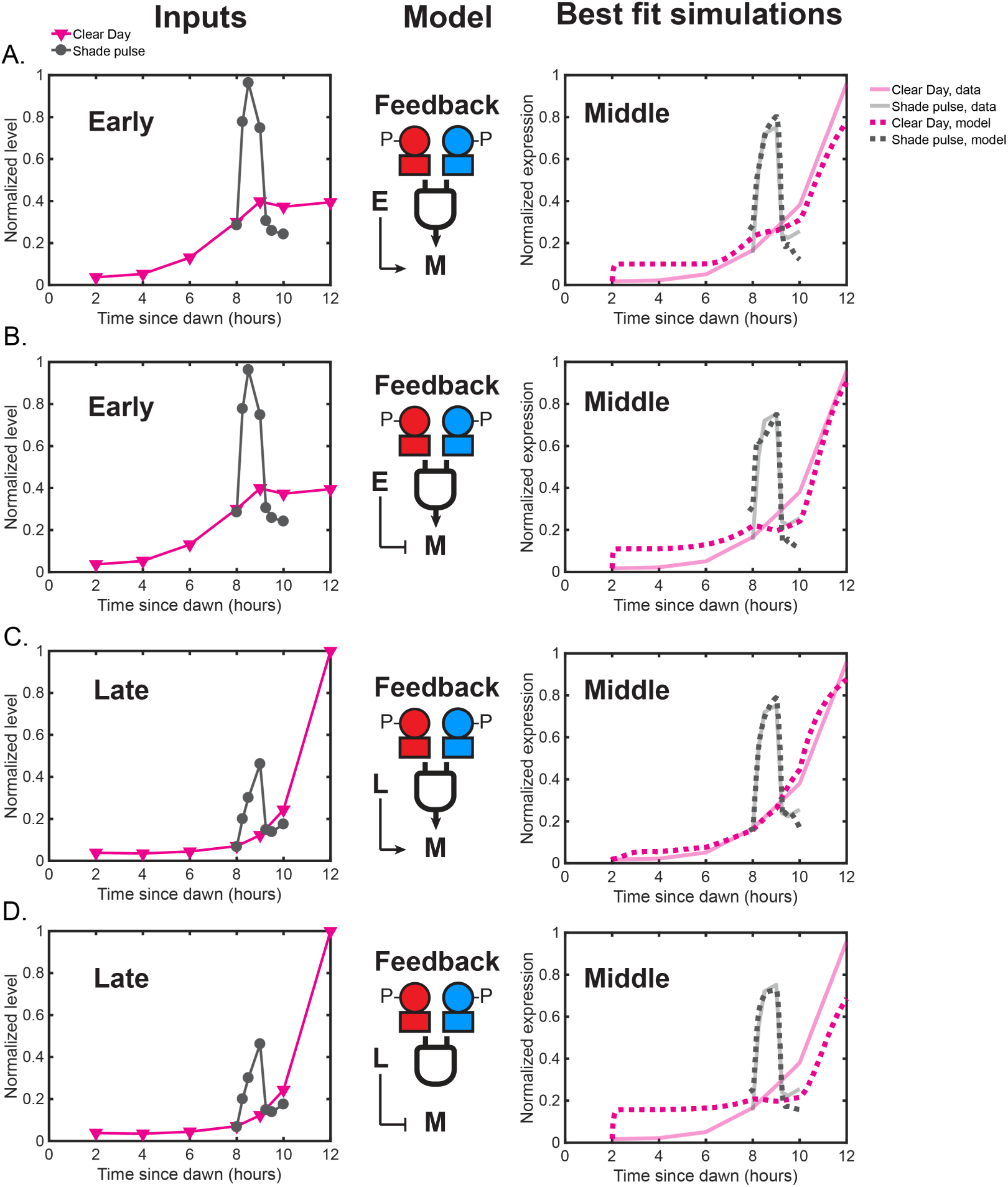
Models in which either the Early or Late cluster feeds back to influence Middle cluster expression. **(A)** Feedback model in which the expression of the Middle dusk cluster is an activation Hill function of Early gene expression and an activation Hill function of both RpaA∼P and RpaB∼P. The left plot shows normalized Early cluster expression levels under Clear Day (magenta) and Shade pulse (gray) conditions used as model input in addition to the RpaA∼P and RpaB∼P dynamics shown in Figure 7B,E of the main text. The right plot shows the average expression values of the Middle cluster data (solid transparent lines), and the simulation produced the best fit model (dotted lines). Data for Clear Day conditions are plotted in magenta, and Shade pulse in gray. **(B)** Feedback model in which Middle gene cluster expression is a repression Hill function of Early cluster expression levels and an activation Hill function of both RpaA∼P and RpaB∼P, presented as in (A). **(C)** Feedback model in which Middle cluster expression is an activation Hill function of Late cluster expression levels and an activation Hill function of both RpaA∼P and RpaB∼P. The left plot shows normalized Late cluster expression levels under Clear Day (magenta) and Shade pulse (gray) conditions used as model input. The right plot shows the average expression values of the Middle cluster data (solid transparent lines), and the simulation produced the best fit model (dotted lines). Data for Clear Day conditions are plotted in magenta, and Shade pulse in gray. **(D)** Feedback model in which Middle cluster expression is a repression Hill function of Late cluster expression levels and an activation Hill function of both RpaA∼P and RpaB∼P, presented as in (C).

**Figure 7-figure supplement 5:**
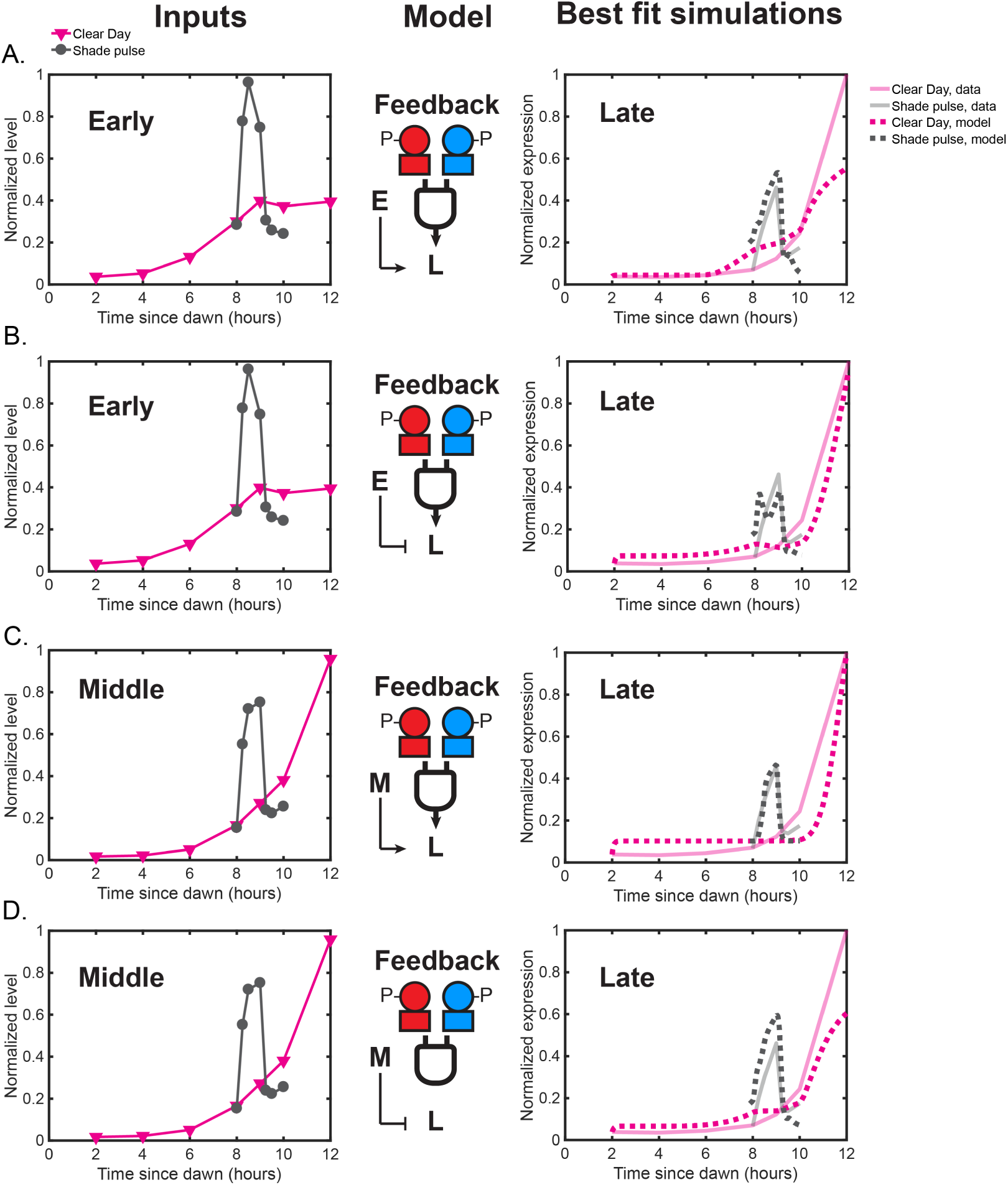
Models in which either the Early or Middle cluster feeds back to influence Late cluster expression. **(A)** Feedback model in which the expression of the Late dusk cluster is an activation Hill function of Early gene expression and an activation Hill function of both RpaA∼P and RpaB∼P. The left plot shows normalized Early cluster expression levels under Clear Day (magenta) and Shade pulse (gray) conditions used as model input in addition to the RpaA∼P and RpaB∼P dynamics shown in Figure 7B,E of the main text. The right plot shows the average expression values of the Middle cluster data (solid transparent lines), and the simulation produced the best fit model (dotted lines). Data for Clear Day conditions are plotted in magenta, and Shade pulse in gray. **(B)** Feedback model in which Late gene cluster expression is a repression Hill function of Early cluster expression levels and an activation Hill function of both RpaA∼P and RpaB∼P, presented as in (A). **(C)** Feedback model in which Late cluster expression is an activation Hill function of Middle cluster expression levels and an activation Hill function of both RpaA∼P and RpaB∼P. The left plot shows normalized Middle cluster expression levels under Clear Day (magenta) and Shade pulse (gray) conditions used as model input. The right plot shows the average expression values of the Late cluster data (solid transparent lines), and the simulation produced the best fit model (dotted lines). Data for Clear Day conditions are plotted in magenta, and Shade pulse in gray. **(D)** Feedback model in which Late cluster expression is a repression Hill function of Middle cluster expression levels and an activation Hill function of both RpaA∼P and RpaB∼P, presented as in (C). A model with an coherent feedforward architecture where the Middle cluster activates the Late cluster (C) best recapitulates the difference of the Late cluster responses to Shade and Clear Day Sunset. During the Shade pulse, the Middle cluster levels do not reach high enough levels to allow for strong activation of Late cluster expression, but at Sunset in Clear Day, Middle cluster levels reach high enough levels to activate the expression of the Late cluster.

